# Protection of photosynthesis by UVR8 and cryptochromes in Arabidopsis under blue and UV radiation

**DOI:** 10.1101/2024.12.31.630900

**Authors:** Luis Orlando Morales, Alexey Shapiguzov, Neha Rai, Pedro José Aphalo, Mikael Brosché

## Abstract

Photosynthesis in plants is negatively affected by high light intensity and UV radiation. The photoreceptors UV RESISTANCE LOCUS 8 (UVR8) and CRYPTOCHROMES (CRYs) mediate perception and acclimation of plants to UV-B/UV-A2 (290–340 nm) and UV-A1/blue light (350–500 nm), respectively. However, their roles in photoprotection of photosynthesis across different wavebands of the spectrum remain unclear. Using chlorophyll fluorescence and LED lighting we studied the roles of UVR8 and CRYs in maintaining photosynthetic capacity in Arabidopsis exposed to UV-B, UV-A1, and blue light. Analysis of quantum yield of Photosystem II, non-photochemical quenching, and LHCII phosphorylation demonstrated that CRYs preserve photosynthetic performance in plants exposed to UV-B, UV-A1, and blue light. UVR8 and CRYs exhibit partially redundant functions in maintaining photosynthetic activity under UV-B, UV-A1, and blue light, and in preventing photodamage under high UV-A1 irradiance. Impaired UVR8 and CRY signaling reduced epidermal flavonol accumulation in leaves, which further compromised photoprotection. These findings provide valuable insights into how UV and blue light perception contribute to photoprotection, with broad implications for plant performance both in natural and managed environments.

## Introduction

UV radiation (UV, 280-400 nm) is an intrinsic component of the electromagnetic spectrum that influences plant performance, affecting metabolism, growth, development, and acclimation of plants to the natural environment (Jenkins, 2009; Barnes *et al*., 2023). UV radiation passing through the stratosphere is spectrally divided into UV-B (280–315 nm; with little radiation below 293 nm reaching the ground) and UV-A (315–400 nm). UV-A can be further divided into UV-A1 (340–400 nm) and UV-A2 (315–340 nm). Plant responses to UV radiation are influenced by the photon energy carried by different wavebands within the UV region, intensity and duration of UV exposure, and by the background photosynthetically active radiation (PAR, 400 – 700 nm) (Jenkins, 2009; Rai, Morales and Aphalo, 2021). Ambient levels of UV activate plant acclimation against potentially harmful, high energetic UV-B radiation. The accumulation of phenolic compounds including flavonoids and phenolic acids, and antioxidants such ascorbic acid, tocopherols, and glutathione as well as changes in amino acid metabolism and hormone signal transduction are well characterized acclimatory responses in plants exposed to UV radiation (Jansen, Gaba and Greenberg, 1998; Demkura *et al*., 2010; Julkunen-Tiitto *et al*., 2014; Vanhaelewyn *et al*., 2016; Podolec, Demarsy and Ulm, 2021; Badmus *et al*., 2022; Palma *et al*., 2022). These UV-mediated changes in metabolism help plants shield cellular components from UV-induced oxidative stress, respond to abiotic and biotic stress and protect key physiological processes including photosynthesis.

Photosynthesis is one of the most crucial biological processes on Earth, yet it remains highly vulnerable to damage in plants subjected to high light intensities and high energy UV radiation (Demarsy, Goldschmidt-Clermont and Ulm, 2017). The impact of UV radiation on photosynthesis is dependent on plant species, UV spectral distribution and irradiance and on the plant physiological status (UV acclimated vs non-acclimated) (Allen, Nogué and Baker, 1998; Jordan, Strid and Wargent, 2016; Sun, Kaiser, Aphalo, et al., 2024). UV-B photons can target several components of the photosynthetic apparatus including both Photosystems (PSII and PSI) where PSII is the most affected (Strid, Chow and Anderson, 1994; Jordan, Strid and Wargent, 2016). UV-B can induce the degradation of the D1 protein and disruption of the manganese cluster of the oxygen evolving complex. It can inhibit the synthesis of photosynthetic pigments including chlorophylls and expression and activities of proteins such as Rubisco, Chl a/b binding proteins (*Lhcb)*, and D1 polypeptide of PSII (*psbA*) (Strid, Chow and Anderson, 1990; Jordan *et al*., 1994; Ranjbarfordoei, Samson and Van Damme, 2011). High levels of UV-A radiation can also target PSII and decrease the maximum quantum efficiency of PSII photochemistry, electron transport rate, and photosynthesis by damaging the oxygen evolving complex and degrading D1 and D2 proteins (Verdaguer *et al*., 2017; Sun, Kaiser, Zhang, *et al*., 2024). At the level of transcription, UV-B radiation impacts on transcript accumulation of photosynthesis related genes (Casati and Walbot, 2003; Rai et al., 2020). To fully understand how UV radiation affects photosynthesis, it is necessary to distinguish between the direct UV damage and the acclimation that mitigates the damage, a task that is not always easy.

Direct damage to photosynthetic machinery can be caused by “excess” photons absorbed by the chloroplasts and insufficient capacity of photosynthetic regulatory mechanisms to quench excessive excitation (Rochaix, 2011). High energy UV photons can also cause damage through additional mechanisms such as disruption of DNA or other macromolecules (Podolec, Demarsy and Ulm, 2021). Accordingly, plants have evolved mechanisms that both prevent and repair damage (Zhang *et al*., 2024). A first line of defence consists in reducing the number of photons reaching the mesophyll and chloroplasts. Physiologically this is achieved through leaf and chloroplast movements or synthesis of photoprotective pigments. The second line of defence uses photosynthetic regulatory mechanisms to dissipate excessive energy of UV photons. These mechanisms include several types of non-photochemical quenching (NPQ), including energy-dependent dissipation of the absorbed light energy in the form of heat (qE), reversible movement of light-harvesting antenna complex II (LHCII) between PSI and PSII called the state transitions (qT), adjustments in the amounts of photosynthetic pigments xanthophylls (qZ) and photoinhibition of PSII (qI) (Nilkens et al., 2010; Shapiguzov and Kangasjärvi, 2022). The third line of defence is to repair damage (Zhang *et al*., 2024). These multiple mechanisms balance efficient capture of photons under lower irradiance with avoidance of damage under exceptionally high irradiance. An important gap in knowledge relates to the role of different photoreceptors in the regulation of the different photosynthesis photo-protection mechanisms conferring tolerance to UV-B and UV-A1 exposure.

Plants use several receptors to perceive light stimuli across different regions of the spectrum. Plant responses to ambient levels of UV-B radiation are largely mediated by the UV photoreceptor UV RESISTANCE LOCUS 8 (UVR8) while those to UV-A/blue radiation (315–500 nm) involve CRYPTOCHROMES 1 and 2 (CRYs) (Ahmad and Cashmore, 1993; Rizzini *et al*., 2011). In sunlight, UVR8 mediates perception of wavelengths shorter than approx. 350 nm (UV-B and UV-A2) and CRYs predominantly mediate responses to longer wavelengths (Rai, Morales and Aphalo, 2021). UVR8 and CRYs are key regulators of gene expression and metabolic responses enabling plants to acclimate and survive under solar UV-B radiation (Rai et al., 2020; Tissot and Ulm, 2020; Rai, Morales and Aphalo, 2021). Plant photoreceptors are not directly involved in photosynthesis; however, their functionality is a prerequisite for normal operation and development of the chloroplast and photosynthetic apparatus (Griffin and Toledo-Ortiz, 2022). Both red- (via phytochromes) and blue light (via CRYs) regulate the transcription of nuclear genes encoding chloroplast localized proteins (Kleine *et al*., 2007). Previous studies using the green alga *Chlamydomonas reinhardtii* and the model plant *Arabidopsis thaliana* showed decreased quantum yield of PSII in *uvr8* mutants exposed to UV-B radiation (Davey *et al*., 2012; Allorent *et al*., 2016; Leonardelli *et al*., 2024). The contrasting levels of sinapate esters in mutants with different levels of UVR8 activity correlate with their sensitivity to UV-B stress and suggests that UVR8 mediates UV-B acclimation and photoprotection through the regulation of phenylpropanoid metabolism (Leonardelli *et al*., 2024). However, as UVR8 and CRYs reciprocally regulate responsiveness to photons (Rai et al., 2019; Tissot and Ulm, 2020), fully understanding of how UVR8 mediates photoprotection requires studies that consider both UVR8 and CRYs.

Even though future climate change scenarios do not anticipate drastic changes in UV-B levels (Barnes *et al*., 2023), solar UV-A radiation at ground level contributes about 95 % of the photons within the UV region and is able to modulate plant metabolism and performance (Morales *et al*., 2010, 2013; Verdaguer *et al*., 2017; Rai, Morales and Aphalo, 2021). As in natural and plant-production environments both UV irradiance and spectral composition vary widely, there is a need to understand how photosynthesis responses to UV radiation of different wavelengths are mediated through coordinated actions of UVR8 and CRYs. In this study, we used light emitting diodes (LEDs) under controlled environment to assess *in vivo* the roles of UVR8 and CRYs in the protection of photosynthetic capacity in plants exposed to UV-B, UV-A1 and blue radiation at irradiances similar to those in sunlight. We show that UVR8 and CRYs control photosynthetic performance by preserving the function of different components of the photosynthetic apparatus. CRYs play major roles in photoprotection in plants exposed to blue light and high UV-A1 irradiance. Furthermore, CRYs and UVR8 play important and partially redundant roles in maintaining photosynthetic activity under UV-B, UV-A1 and blue light and in the avoidance of photodamage under high UV-A1 irradiance.

## Materials and Methods

### Plant material, growth, and light treatments

Arabidopsis accessions Landsberg erecta (L*er*) and Wassilewskija (Ws) were used in the experiments. Photoreceptor mutants including *uvr8-2* (Brown *et al*., 2005), *cry1cry2* (Mazzella *et al*., 2001) and *uvr8-2cry1cry2* (Rai *et al*., 2019) are in L*er* background while *uvr8-7* and the overexpression line *UVR8*-OE (Favory *et al*., 2009) are in Ws. Seeds were sown in plastic pots (8 cm × 8 cm) containing a 1:1 mixture of peat and vermiculite and kept in darkness at 4°C for 3 days. Subsequently, pots were transferred to a controlled environment growth room at 23 °C:19°C and 70%:90% relative humidity (light:dark) under 12 h photoperiod (07:00 to 19:00) from fluorescent tubes (Osram T8 L 36 W/865 Lumilux) with photon irradiances of 220 µmol m^-2^ s^-1^ PAR, < 0.12 µmol m^-2^ s^-1^ of UV-B radiation and 0.32 µmol m^-2^ s^-1^ of UV-A2 and 1.47 µmol m^-2^ s^-1^ of UV-A1 radiation (Fig. S1), where they remained for 7 days. Thereafter, plants were transplanted to a tray containing six 4 x 4 cm pots where seedlings of the different genotypes were combined. Plants were further grown for 10 days in the growth room under the same conditions.

When plants were 17 days old, they were exposed to different light treatments lasting for 20 h or 90 min (Table 1). For this, plants were moved from the growth rooms to the light treatments in the afternoon between 14:00 and 16:00. The treatments were created using different LEDs driven at a set constant current: UV-B (313 nm, UVMAX305, Roithner Lasertechnik GmbH, Austria), Blue and Blue + Blue (448 nm, actinic light provided by the Imaging PAM, M-series; Walz, Germany) and UV-A1 (369 nm, LZ1-10UV00, LED Engin, San Jose, CA). Table 1 and Fig. S1 show the photon irradiance and spectral distribution for every light treatment used. The irradiance selected for each waveband is comparable to clear-sky sunlight levels during summertime in Helsinki, Finland. The spectral irradiance was measured with an array spectroradiometer (Maya 2000Pro, Ocean Optics Inc., Dunedin, FL) using R (R Core Team, 2023) with packages ‘photobiology’, ‘photobiology Wavebands’ and ‘ooacquire’ from the R for Photobiology suite (Aphalo, 2015).

**Table 1.** Light treatments used in the experiments. The treatments were single irradiation events of the duration indicated. Some treatments consisted in light from a one nearly monochromatic source and others in the combined light from two such sources. Light emitting diodes (LEDs) were used, as described in the text.

|  | Treatments |  |  |  |  |  |  |
| --- | --- | --- | --- | --- | --- | --- | --- |
|  | Blue | Blue + Blue | UV-A1 | UV-A1 + Blue | UV-B | UV-B + Blue | UV-A1 *short* |
|  | Duration of treatments (h) |  |  |  |  |  |  |
|  | 20 | 20 | 20 | 20 | 20 | 20 | 1.5 |
| | Photon irradiance ($\mu\text{mol m}^{-2} \text{s}^{-1}$) | | | | | | |
| Blue | 204 | 286 | 0 | 204 | 0 | 193 | 0 |
| UV-A1 | 0 | 0 | 100 | 95.6 | 0 | 0 | 100 |
| UV-B | 0 | 0 | 0 | 0 | 1.1 | 1.56 | 0 |

### Chlorophyll fluorescence measurements

Multiple photosynthetic parameters were assessed by room-temperature chlorophyll fluorescence imaging, which allowed simultaneous measurements of multiple rosettes of each of the different genotypes studied. Concurrently with the 20 h-long light treatments described above; the photosynthetic parameters were measured repeatedly to assess the time course of response. Trays with six potted plants per genotype were placed under the Imaging PAM and images of chlorophyll fluorescence were recorded as described in (Wang *et al*., 2020) with minor modifications. At the beginning of each experiment, minimal (Fo) and maximal (Fm) fluorescence were determined in plants that were dark-acclimated for 30 minutes. Subsequently, light treatments were started with continuous monitoring of steady-state fluorescence (Fs; every 2 min) and maximal light-acclimated fluorescence (Fm′; every 10 min). The effective quantum yield of PSII photochemistry (φPSII) was calculated as φPSII = (Fm′ – Fs)/Fm′ (Genty, Briantais and Baker, 1989). Non-photochemical quenching (NPQ) was calculated after 30 min of light treatment as (Fm-Fm’)/Fm’ (Horton and Ruban, 1992).

To assess UV-A1-induced photoinhibition, non-treated plants from the growth rooms were acclimated to darkness for at least 30 min and exposed to UV-A1 for 90 min, then UV-A1 was switched off and recovery was followed in darkness with Fs and Fm measurements as described above. Photoinhibition in presence of lincomycin (2 mM) or methyl viologen (0.15 µM) was assayed with the Imaging PAM in leaf discs, after overnight acclimation in darkness in the presence of the chemicals, as described in (Shapiguzov and Kangasjärvi, 2022).

### Biochemical analyses of photosynthesis

To address the effects of different 20-h light treatments on the photosynthetic apparatus at the biochemical level, six rosettes of each genotype exposed to each of the light treatments were snap-frozen in liquid nitrogen and stored at –80°C at the end of the light exposure. Total protein was extracted, separated by SDS-PAGE and immunostained with antibodies against the D1 subunit of PSII (Agrisera) and with phospho-threonine/tyrosine antibody (Cell Signaling) to assess phosphorylation of PSII light-harvesting antennae (LHCII) (Shapiguzov *et al*., 2010).

### Non-destructive Optical Absorbance Measurements

To assess how the 20 h UV-B, UV-A1, Blue and combined light treatments affected flavonol, chlorophyll and anthocyanin contents in different genotypes, non-destructive measurements of epidermal absorbance were performed with a Dualex Scientific+ device (Force-A™, France). The UV-A1 (375 nm) absorbance of the adaxial epidermis was assessed in four developed leaves per plant from at least five plants per genotype and treatment. Plants that remained in the growth room and those treated were measured in random sequence right after the 20 h treatments were finished. The measurements were performed in four independent experiments.

### Statistical analysis

Significant effects (*P < 0.05*) of light treatments on photosynthetic parameters were assessed using One-Way Analysis of Variance (ANOVA) followed by post-hoc Tukey test in IBM SPSS Statistics. Dualex and NPQ data for each light treatment were analyzed separately, using Linear mixed-effects models with replicate runs of experiments as grouping factors and a random effect for the intercept using the R package NLME (Pinheiro and Bates, 2024). For these data, ANOVA was used to assess significant effects (*P < 0.05*) of the genotype on flavanol, chlorophyll and anthocyanin content under individual light treatments. Comparisons between genotypes were assessed by fitting contrasts using the R package gmodels 2.18.1 (Gregory Warnes *et al*., 2018). Figures were plotted using the R package ggplot2 (Wickham, 2009) version 3.4.2 and Microsoft Excel.

## Results

### UVR8 and CRY signaling promote favourable adjustment of PSII functioning under UV and blue radiation

We evaluated the contribution of photoreceptors to the regulation of PSII activity under different light treatments that activate UVR8 (UV-B, 310 nm) and CRY signaling through UV-A1 (365 nm) and blue light (450 nm) (Rai et al., 2020). The UV-B and UV-A1 treatments were applied alone or in combination with blue light to assess possible effects of crosstalk between UVR8 and CRY signaling. Quantum yield of PSII photochemistry (φPSII) was continuously measured during 20 h under an Imaging PAM in L*er*, *uvr8-2*, *cry1cry2* and *uvr8-2cry1cry2* Arabidopsis plants grown for 17 days under white light and then subjected to different light treatments (Fig. 1).

**Figure 1.**
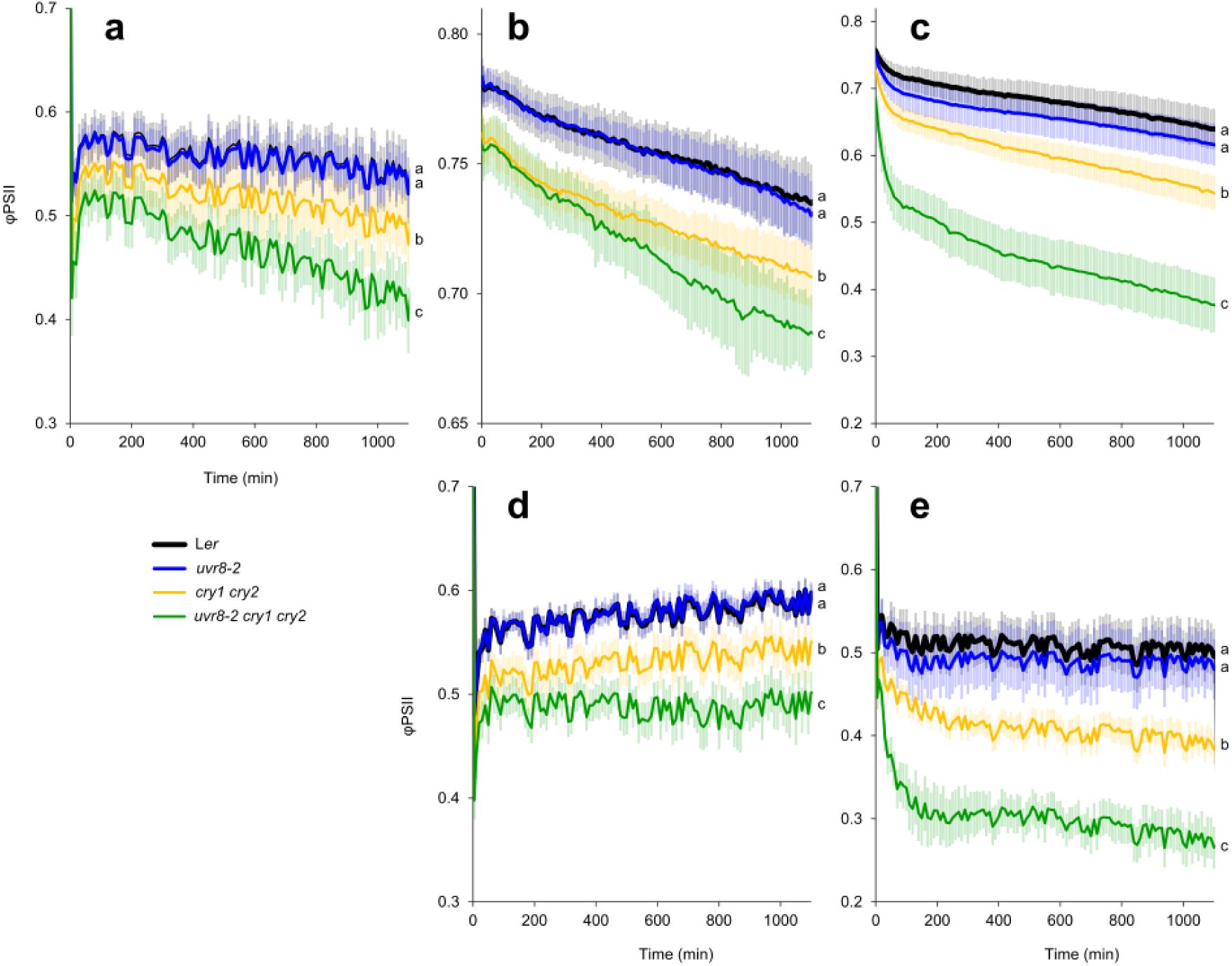
Quantum yield of PSII photochemistry (φPSII) measured in L*er*, *uvr8-2*, *cry1cry2* and *uvr8-2cry1cry2* with Imaging PAM during 20 h under different light treatments. a (Blue light, 448 nm 204 µmol m-2 s-1), b (UV-B, 313 nm 1 µmol m-2 s-1), c (UV-A1, 369 nm 100 µmol m-2 s-1), d (UV-B + Blue) and e (UV-A1 + Blue). The data points represent means, and the error bars indicate the SD. For each experiment at least 6 individual rosettes per genotype were measured. The experiment was repeated at least three times with similar results. Significant differences (*P<0.05*) between genotypes after 20 h of exposure to each light treatment are denoted with different letters. Be aware that the y-axis scale differs among panels.

Exposure of plants to blue light (204 µmol m^-2^ s^-1^) produced a small but significant drop in φPSII in the *cry1cry2* mutant compared to L*er* (Fig. 1a). The drop in φPSII was larger in the triple mutant than in *cry1cry2* and *uvr8-2* while *uvr8-2* and L*er* had similar φPSII (Fig. 1a, Table S1). Increasing blue light irradiance (Blue + Blue, 286 µmol m^-2^ s^-1^) showed the same φPSII responses in the four genotypes compared to 204 µmol m^-2^ s^-1^ (Fig. S2). When plants were exposed to UV-B and UV-A1, a similar response between genotypes was observed to that of blue light (Figs 1b, c). The *uvr8-2cry1cry2* and *cry1cry2* showed a significant decrease in φPSII compared to L*er* and *uvr8-2*. However, PSII damage in *uvr8-2cry1cry2* appeared to be worse and it progressed faster under UV-A1 than blue light or UV-B (Figs 1a-c). Exposure to UV-B and UV-A1 together with blue light affected the dynamics of φPSII but did not change the relative genotype responses compared to UV-B or UV-A1 alone (Figs 1d, e). Taken together our data indicate an important role of CRY signaling on PSII photochemistry under blue light, UV-B and UV-A1. Furthermore, simultaneously impaired function of CRY and UVR8 signaling was more detrimental for plant photosynthesis than the depletion of CRYs alone.

A decrease of φPSII during light exposure is caused by non-photochemical quenching (NPQ) (Baker, 2008). NPQ consists of several components including energy-dependent qE, qZ, qT and qI (Nilkens et al., 2010; Shapiguzov and Kangasjärvi, 2022). As expected, the results in Fig 1 revealed formation of NPQ already in the first minutes of exposure to the light treatments. To compare the effects of the treatments in different genotypes, we measured NPQ after 30 min of light exposure (Fig. S3). At this early time point we expect NPQ to mostly consist of qE and qT (but not qZ or qI). NPQ was similar in the tested genotypes, except for UV-A1. Under UV-A1 alone or together with blue, the *cry1cry2* and *uvr8-2cry1cry2* mutants developed significantly higher NPQ than L*er*. In contrast, no differences in NPQ between the genotypes were observed after a 30-min UV-B exposure. Interestingly, UV-B alone caused negative apparent NPQ values in all genotypes likely because Fm’ under UV-B was higher than the dark-adapted Fm. This can be explained by photosynthetic state transitions (qT), the relocations of light harvesting antenna complex II (LHCII) between PSI and PSII. We hypothesize that UV-B triggered migration of the mobile pool of LHCII to PSII (formation of state 1), which increased light-absorption cross-section of PSII and thus Fm’ (see also Fig. 6).

To further study the possible role of UVR8 in blue light, UV-B and UV-A1 responses related to photosynthesis, we performed similar analyses in the UVR8 overexpressor line (*UVR8*-OE), *uvr8-7* knockout mutant and their corresponding wildtype accession, Ws (Fig. 2). Like in the case of L*er* and *uvr8-2,* φPSII was affected in the same way in *uvr8-7* and Ws under all light treatments (Figs 2 a-e, Table S1). However, in *UVR8*-OE the tolerance of φPSII to blue light, UV-B and UV-A1 radiation was significantly higher than in Ws and *uvr8-7* (Figs 2 a-e), supporting UVR8 functions in UV-B and involvement in UV-A1 and blue light responses (Rai et al., 2020). Exposure to UV-B and UV-A1 together with blue light did not change the relative genotype responses compared to UV-B or UV-A1 alone (Figs 2 a-e).

**Figure 2.**
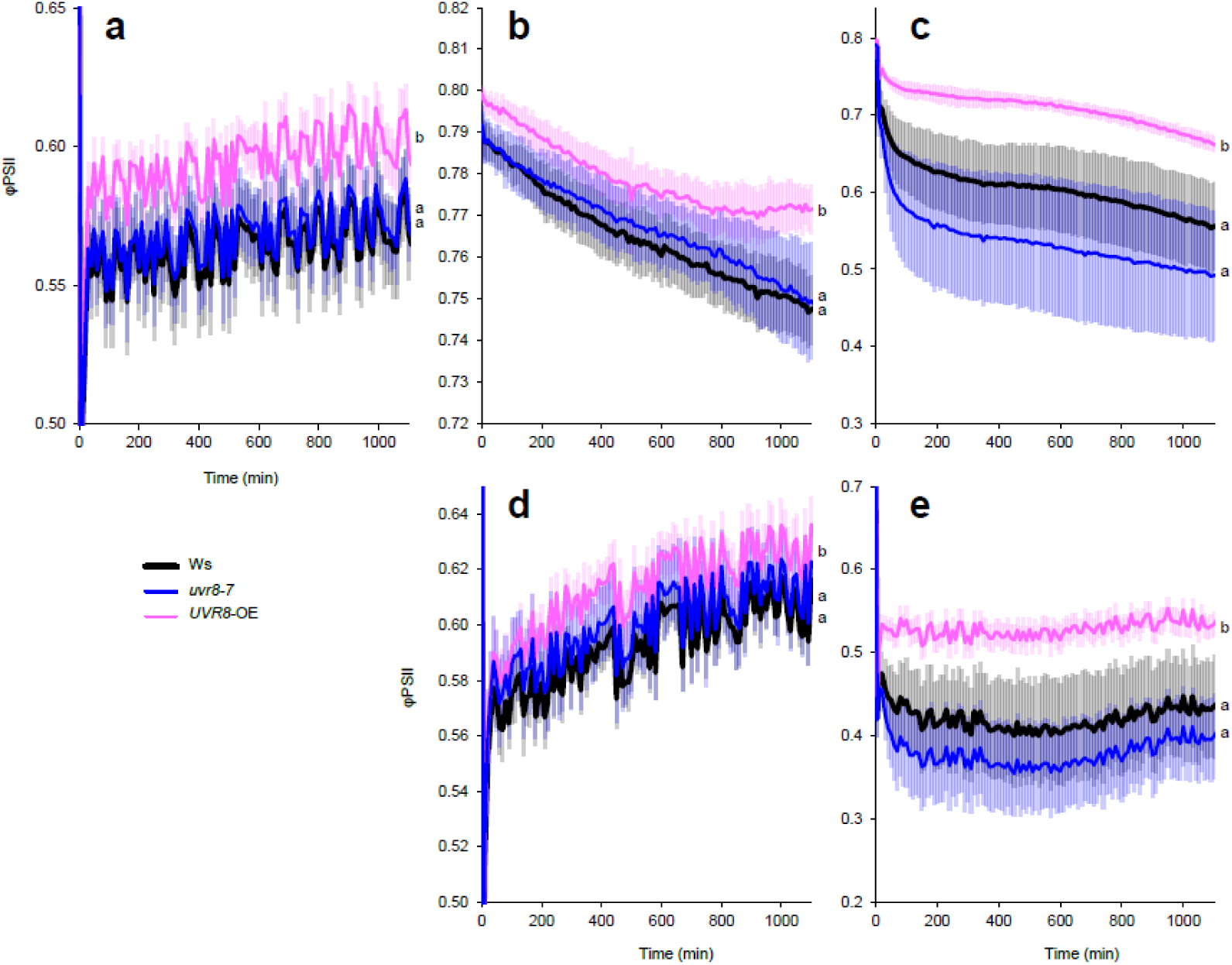
Quantum yield of PSII photochemistry (φPSII) measured in wildtype Ws, *uvr8-7* and *UVR8*-OE with Imaging PAM during 20 h under different light treatments. a (Blue light, 448 nm 204 µmol m^-2^ s^-1^), b (UV-B, 313 nm 1 µmol m^-2^ s^-1^), c (UV-A1, 369 nm 100 µmol m^-2^ s^-1^), d (UV-B + Blue) and e (UV-A1 + Blue). The data points represent means, and the error bars indicate the SD. For each experiment at least 6 individual rosettes per genotype were measured. The experiment was repeated at least three times with similar results. Significant differences (*P<0.05*) between genotypes under each light treatment are denoted with different letters. Be aware that the y-axis scale differs among panels.

### CRYs and UVR8 protect Arabidopsis from PSII photoinhibition

The components of NPQ (qE, qZ, qT and qI) have different relaxation kinetics in darkness, with qI being the slowest (Shapiguzov and Kangasjärvi, 2022). After 20 h light exposure (Figs 1, 2), plants were acclimated to darkness for at least 30 min, which is sufficient to relax qE, and measured dark-adapted φPSII (Fv/Fm). We observed that blue light (204 µmol m^-2^ s^-1^) induced photoinhibition to the same extent in *cry1cry2* and *uvr8-2cry1cry2* compared to L*er* and *uvr8-2*, while increasing blue light irradiance to 286 µmol m^-2^ s^-1^ (Blue + Blue) increased photoinhibition further in the triple mutant compared to *cry1cry2* (Fig. 3 a, Table S1).

**Figure 3.**
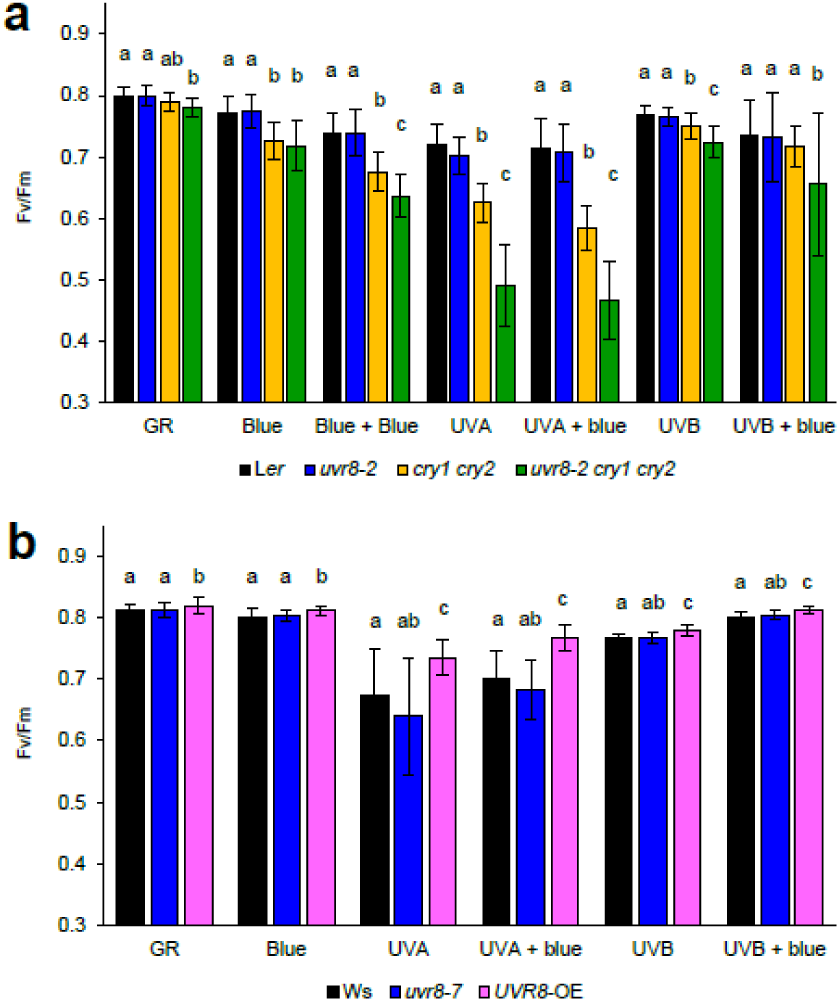
Fv/Fm measured in dark-adapted plants that were exposed to different light treatments for 20 h. Blue (Blue light, 448 nm 230 µmol m^-2^ s^-1^), Blue + Blue (Blue light, 448 nm 286 µmol m^-2^ s^-1^), UVA (UV-A1, 369 nm 100 µmol m^-2^ s^-1^), UVB (UV-B, 313 nm 1 µmol m^-2^ s^-1^), GR (plants grown in growth rooms in parallel with treated plants - but not treated with UV or blue). (A) Photoreceptor mutants in L*er* background and (B) in Ws background. The bars represent means of three independent biological repeats, and the error bars indicate the SD. In each experiment n = 6 plants of each genotype were measured. Significant differences (*P<0.05*) between genotypes under each light treatment are denoted with different letters.

The UV-B treatment lowered Fv/Fm values in *cry1cry2* compared to L*er* and to even lower levels in the triple mutant compared to *cry1cry2*; however, UV-B exposure in the presence of blue light resulted in no significant differences between Fv/Fm values recorded for L*er* and *cry1cry2* (Fig. 3a). Furthermore, in agreement with the data presented in Fig. 1, the UV-A1 treatment revealed persistent and pronounced UV-A1-induced photoinhibition in *uvr8-2cry1cry2* that was significantly different from all genotypes both under UV-A1 alone and in the UV-A1 and blue light treatment (Fig. 3a). Measurements of plants grown in the growth room without exposure to any light treatments showed slightly lower Fv/Fm values in the triple mutant that were significantly different from L*er* and *uvr8-2* (Fig. 3a). In the same assay, under all light treatments *UVR8*-OE was significantly more tolerant to photoinhibition than the wildtype Ws, but the effect was large only under UV-A1 alone or together with blue light (Fig. 3b, Table S1).

To get further insight into the observed decrease in PSII photochemistry of *uvr8-2cry1cry2* and *cry1cry2* under high UV-A1 irradiance, we subjected the plants to a shorter UV-A1 treatment (1.5 h) (Fig. 4).

**Figure 4.**
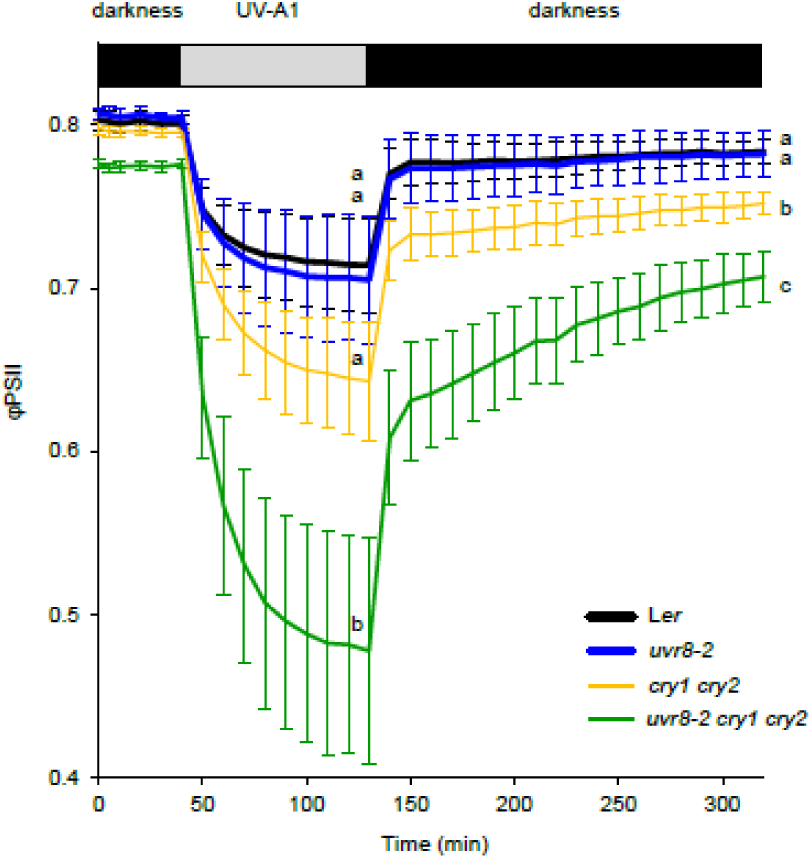
Measurements of φPSII during 1.5 h of UV-A1 treatment (369 nm, 100 µmol m^-2^ s^-1^) and subsequent dark recovery. The data points represent means of three independent biological repeats, and the error bars indicate the SD. In each experiment n = 6 plants of each genotype were measured. Significant differences (*P<0.05*) between genotypes are denoted with different letters.

After this, UV-A1 was turned off and recovery of φPSII was monitored in darkness, with saturating light pulses triggered in 10-minute intervals to assess φPSII. We observed similar φPSII response in L*er* and *uvr8-2* after 1.5h of UV-A1 exposure while a larger but non-significant decreased in φPSII was observed in *cry1cry2*. However, the triple mutant demonstrated much lower φPSII compared to the other genotypes (Fig. 4, Table S1). After the onset of darkness, φPSII quickly recovered and reached a plateau in L*er* and *uvr8-2*, the recovery was incomplete in *cry1cry2* compared to L*er* and incomplete and slower in *uvr8-2cry1cry2* (Fig. 4). This suggested that UV-A1-dependent decrease of φPSII in *uvr8-2cry1cry2* was to a large extent due to qI, and possibly qZ. Before treatment, the variation among plants was very small, and it increased drastically during the first half of the UV-A1 irradiation period. During recovery in darkness, variation also decreased but remained higher than before irradiation. φPSII did not fully recover to the pre-irradiation values in any of the genotypes, but the decrease in φPSII was very small in L*er* and *uvr8-2* (Fig. 4).

To find out whether the high predisposition of *uvr8-2cry1cry2* to PSII photoinhibition was specific to light treatments, we tested tolerance of the wildtype and mutants to other qI-inducing treatments (Fig. 5).

**Figure 5.**
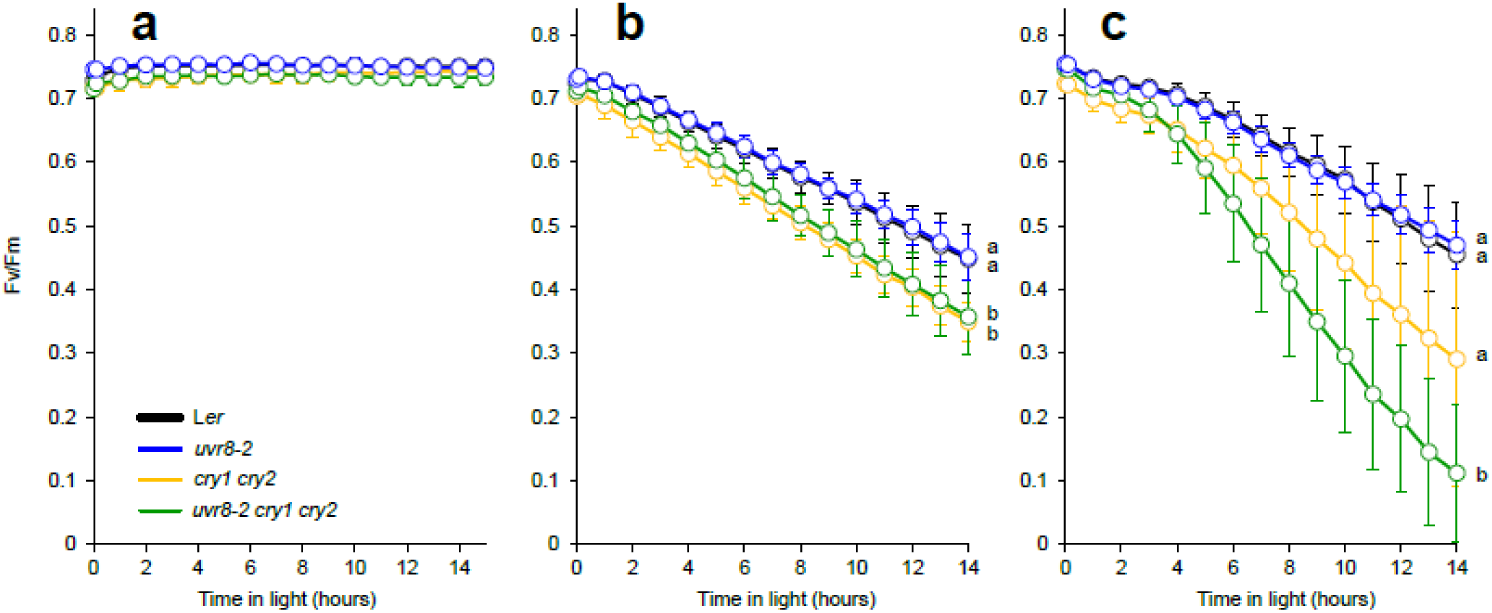
Fv/Fm measured in leaf discs of L*er*, *uvr8-2*, *cry1cry2* and *uvr8-2cry1cry2* exposed to (a) Blue light (448nm, 80 µmol m-2 s-1), (b) Lincomycin and (c) Methyl viologen. The data points represent means of 6 rosettes per genotype and the error bars indicate the SD. The experiment was repeated at least three times with similar results. Significant differences (*P<0.05*) between genotypes are denoted with different letters.

For this, we pre-treated leaf discs with the chemicals lincomycin, an inhibitor of organelle translation and PSII repair; or methyl viologen (MV), which produces reactive oxygen species (ROS) in the chloroplast at PSI. We then exposed leaf discs to repeated 1-hour cycles of blue light (450 nm 80 µmol m^-2^ s^-1^) inside the Imaging PAM. After each light cycle, 23-min dark acclimation was introduced followed by a saturating light pulse to measure Fv/Fm (Fig. 5). In the control treatment without the chemicals, Fv/Fm did not differ between genotypes (Fig. 5a). However, the treatment with lincomycin reduced Fv/Fm values over the time course in all genotypes with end values significantly lower in *cry1cry2* and the triple mutant compared to L*er* and *uvr8-2* (Fig. 5b, Table S1). In response to the ROS-generating compound MV, the triple mutant showed dramatic and significant decrease of Fv/Fm indicating stronger MV sensitivity compared to all genotypes (Fig. 5c). The *uvr8-2* mutant was indistinguishable from L*er* (Figs 5b, c). We also detected higher tolerance to MV in *UVR8*-OE than in Ws (Fig. S4). Taken together our data indicate that CRY signaling and coordination between UVR8 and CRYs is required to maintain PSII photochemistry (φPSII) under conditions leading to the generation of ROS or when PSII repair is compromised.

### Analyses of LHCII phosphorylation

To assess the effects of UV-A and UV-B radiation on the functioning of PSI, PSII and the redox state of the electron-transport chain at the biochemical level, we extracted total protein and assayed by immunoblotting the levels of D1 protein, LHCII and phosphorylated LHCII (P-LHCII) at the end of the 20 h treatments described above (Fig. 6, S5).

**Figure 6.**
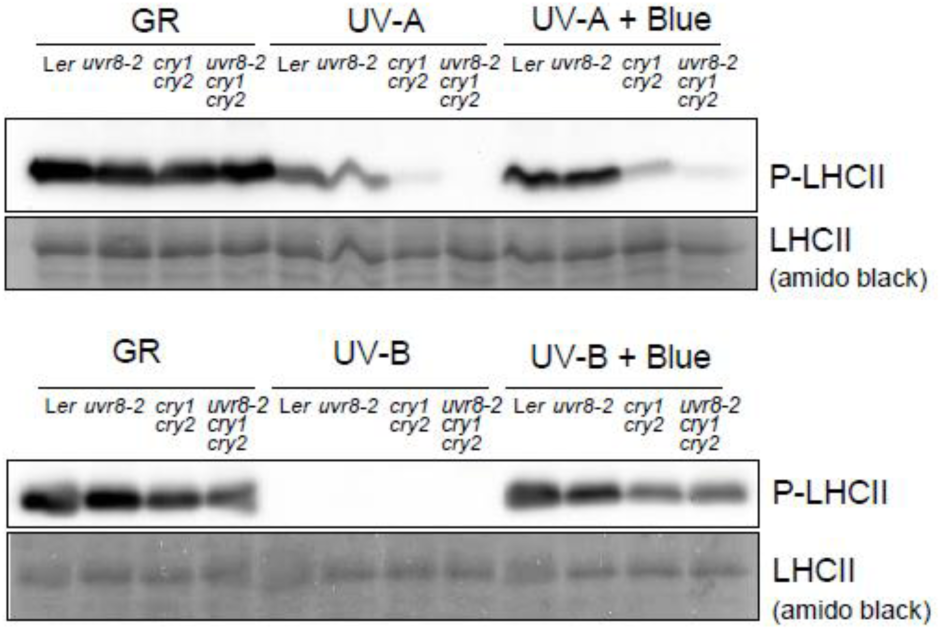
Phosphorylation patterns of LHCII measured in L*er*, *uvr8-2*, *cry1cry2* and *uvr8-2cry1cry2* exposed to UVA1 (369 nm 100 µmol m^-2^ s^-1^), UV-A1 + Blue light, 448 nm 204 µmol m^-2^ s^-1^, UV-B (313 nm 1 µmol m^-2^ s^-1^), and UV-B + Blue light, 450 nm 230 µmol m^-2^ s^-1^, or GR (plants grown in growth rooms in parallel with treated plants - but not treated with UV or blue). The experiment was repeated two independent times.

The phosphorylation of LHCII is sensitive to redox state of photosynthetic electron transport chain between PSII and PSI. LHCII phosphorylation usually increases when PSII is more active than PSI and decreases when PSI is more active than PSII. The levels of P-LHCII differed dramatically between the genotypes (Fig. 6). Under UV-A1, L*er* maintained its levels of P-LHCII while a slight reduction of LHCII phosphorylation was observed in *uvr8-2* (Fig. 6a). The *cry1cry2* mutant showed a drastic reduction in P-LHCII compared to L*er* and *uvr8-2*. Furthermore, there was no detectable LHCII phosphorylation in *uvr8-2cry1cry2* (Fig. 6a). It is worth noting that no changes in total LHCII abundance (amido black staining) were detected under either light treatment. These results agreed well with the deterioration of PSII function in the same mutants observed by chlorophyll fluorescence imaging (Figs 1, 3, 4). Similarly to the chlorophyll fluorescence assays, the most pronounced differences between the genotypes were observed in response to UV-A1. Importantly, no significant decrease was observed in the total levels of PSII under UV-A1 (anti-D1 antiserum) (Fig. S5). This indicated that the observed responses under high UV-A1 irradiance occur on the level of function and not abundance of PSII and LHCII.

LHCII phosphorylation was completely missing in plants exposed to UV-B alone for 20 h, but not under UV-B in the presence of blue light (Fig. 6b). This was likely explained by qT and was in accordance with the chlorophyll fluorescence analyses of NPQ (Fig. S3). The *cry1cry2* and *uvr8-2cry1cry2* mutants showed a mild reduction in P-LHCII compared to L*er* and *uvr8-2* under UV-B plus blue light (Fig. 6b). Taken together, high UV-A1 irradiance had a larger negative impact than low levels of UV-B on φPSII or Fv/Fm (Figs 1, 3, 4) and on PSII biochemistry (Fig. 6).

### UVR8 and CRY-mediated accumulation of epidermal flavonols, chlorophyll and anthocyanins

Flavonols are phenolic compounds induced by UV radiation and high light which help protect the plant from UV and oxidative stress (Hideg, Jansen and Strid, 2013). When they accumulate in the epidermis they attenuate UV-A- and to a lesser extent UV-B radiation before they reach the mesophyll. To test whether the differences we observed in photoinhibition and photosynthetic performance between the studied genotypes could be linked to this protection mechanism, we assessed the epidermal flavonol content after the 20 h light treatments in photoreceptor mutants of both L*er* and Ws backgrounds (Figs 7, 8).

**Figure 7.**
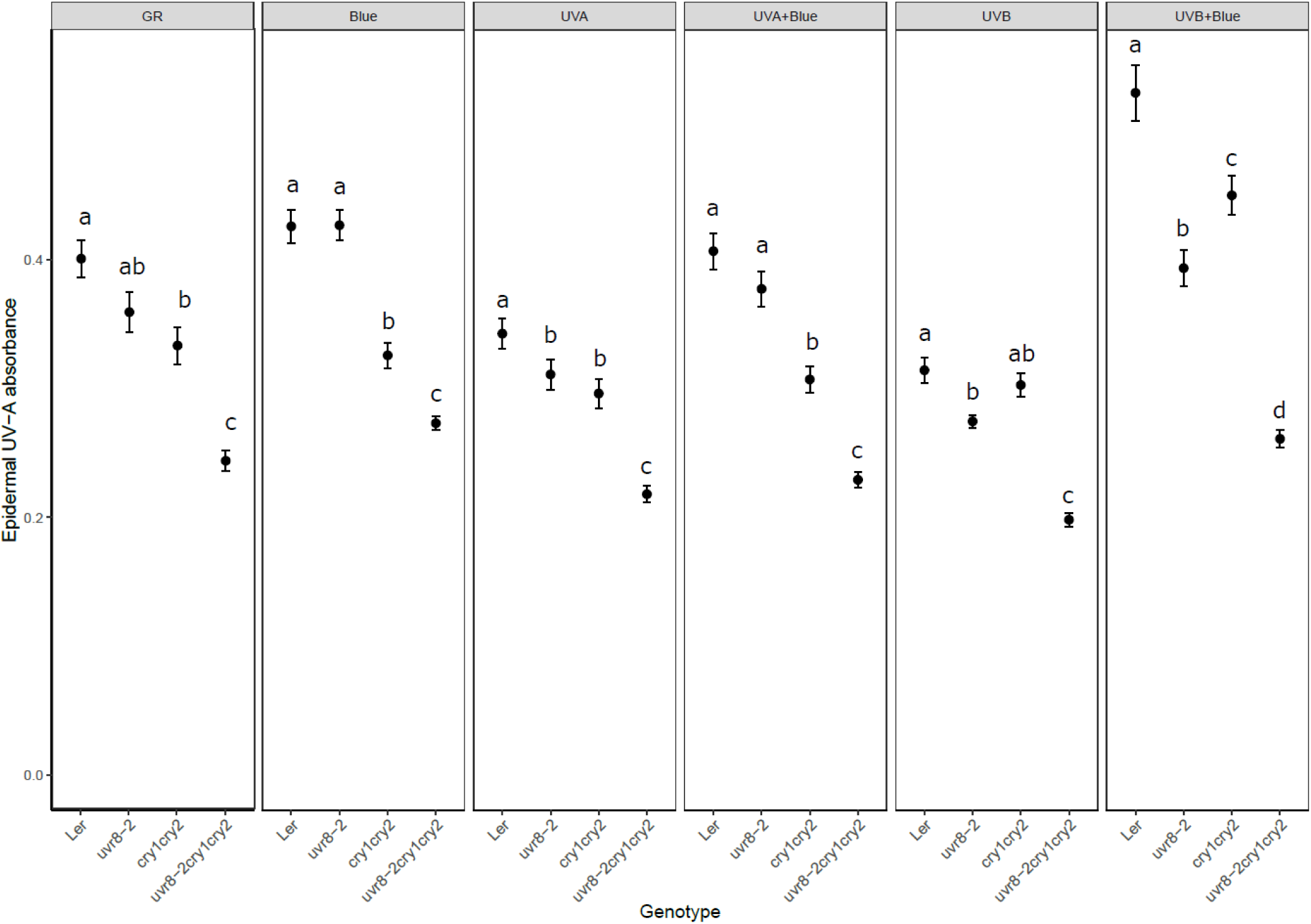
Flavonol content estimated with Dualex in L*er* background genotypes exposed to different light treatments for 20 h. Blue (Blue light, 448 nm 230 µmol m^-2^ s^-1^), UVA (UV-A1, 369 nm 100 µmol m^-2^ s^-1^), UVB (UV-B, 313 nm 1 µmol m^-2^ s^-1^), GR (plants grown in growth rooms in parallel with treated plants - but not treated with UV or blue). The data points represent means of four independent biological repeats, and the error bars indicate the SE. In each experiment two leaves from 6 plants of each genotype were measured. Significant differences (*P<0.05*) between genotypes under each light treatment are denoted with different letters.

The *cry1cry2* mutant showed lower epidermal UV-A1 absorbance than wildtype L*er* under growth room conditions (GR), and after exposure to blue light, UV-A1, UV-A1 plus blue light and UV-B plus blue light (Fig. 7). This indicated lower levels of flavanols in the double *cry* mutant across most studied light treatments. A significantly lower epidermal flavanol content was observed in leaves of *uvr8-2* compared to L*er* in plants exposed to UV-A1, UV-B and UV-B in the presence of blue light but not under blue light alone. In agreement with the previous analyses of photosynthesis, the triple mutant showed the weakest epidermal absorbance of all genotypes under all treatments (Fig. 7, Table S1). We also detected lower levels of epidermal flavonols in *uvr8-7* as compared to Ws under blue light, UV-A1 plus blue, UV-B and UV-B plus blue light (Fig. 8). *UVR8*-OE had significantly higher levels of flavonols than Ws and *uvr8-7* under all light treatments (Fig. 8). The flavonols contents of Ws, *uvr8-7* and *UVR8*-OE were also affected differently under GR conditions (Fig. 8). Taken together, UVR8 and CRY signaling regulate flavonol accumulation under light conditions where these photoreceptors are activated, with the strongest UV-A1 attenuation in the adaxial epidermis of the wildtypes, L*er* and Ws, under simultaneous exposure to both UV-B and blue light.

**Figure 8.**
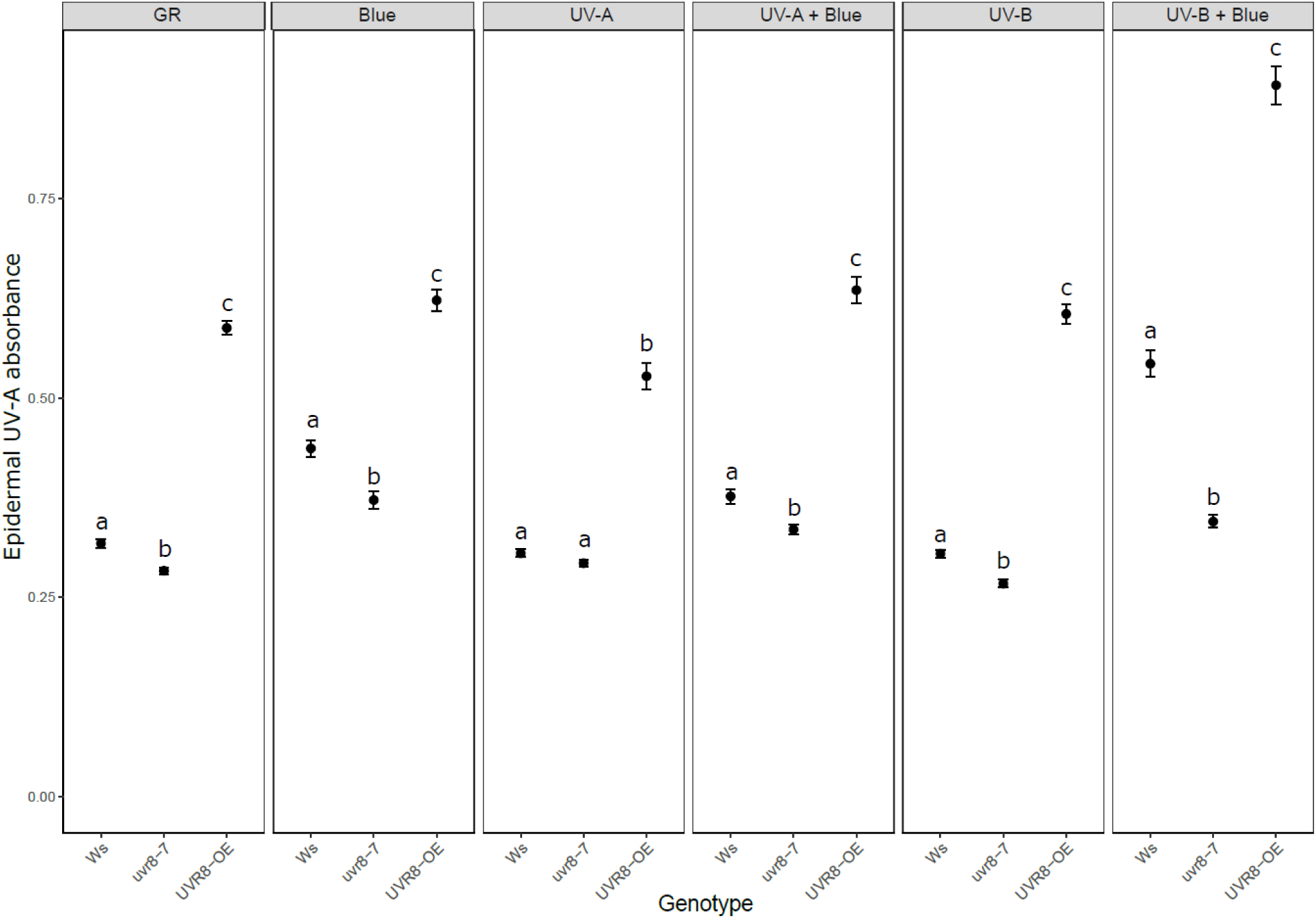
Flavonol content estimated with Dualex in Ws background genotypes exposed to different light treatments for 20 h. Blue (Blue light, 448 nm 230 µmol m^-2^ s^-1^), UVA (UV-A1, 369 nm 100 µmol m^-2^ s^-1^), UVB (UV-B, 313 nm 1 µmol m^-2^ s^-1^), GR (plants grown in growth rooms in parallel with treated plants - but not treated with blue or UV). The data points represent means of four independent biological repeats, and the error bars indicate the SE. In each experiment two leaves from 6 plants of each genotype were measured. Significant differences (*P<0.05*) between genotypes under each light treatment are denoted with different letters.

The optical assessment of chlorophyll can reveal the ability of a leaf to absorb red photons, which indirectly provides an estimate of chlorophyll concentration. The double *cry* mutations had a significant impact on the estimated chlorophyll content under different light treatments (Fig, S6). Compared to L*er*, *cry1cry2* plants had lower levels of chlorophyll under GR conditions, blue light and UV-B radiation (Fig. S6). Chlorophyll contents in *uvr8-2* was increased by UV-A1 in the presence of blue light and reduced under UV-B as compared to L*er* (Fig. S6). Like their flavonol profiles (Fig. 7), triple mutant plants showed significantly lower chlorophyll levels than all the other genotypes under all light conditions tested (Fig. S6). The anthocyanin content was higher in *cry1cry2* than in L*er* and *uvr8-2* under GR, UV-A1 and UV-A1 in the presence of blue light while the single *uvr8-2* mutation had no effects on the anthocyanin levels (Fig. S7). However, simultaneous inactivation of UVR8 and CRYs in the triple mutant resulted in significantly higher anthocyanin levels under GR, blue light, UV-A1, UV-A1 in the presence of blue light and UV-B compared to L*er*, *uvr8-2* and *cry1cry2* (Fig S7).

UVR8 activity modulated chlorophyll and anthocyanin contents in GR grown plants and in those exposed to UV-B and UV-B in the presence of blue light (Figs S8, 8). *UVR8*-OE plants show higher chlorophyll but lower anthocyanin levels than Ws and *uvr8-7* under these treatments (Figs S8, 8). Overall, we observed good connection between the ability of the studied genotypes to accumulate flavonoids during the light treatment with resistance of their photosynthetic apparatus against photoinhibition.

## Discussion

Plants have developed multiple mechanisms to protect the photosynthetic apparatus from potentially harmful radiation while harvesting light energy for photosynthesis. Here, we revealed key roles for UVR8 and CRYs in regulating different aspects of photosynthesis in Arabidopsis by using narrowband irradiation treatments covering UV-B, UV-A1 and blue radiation. We demonstrate that CRYs maintain photosynthetic activity under light conditions where they are activated (UV-A1 and blue light). Furthermore, UVR8 and CRYs redundantly promote normal performance of PSII in response to blue light, UV-B and UV-A1 and protect plants from photodamage induced by high UV-A1 irradiance. Our study reveals wavelength-specific and photoreceptor-dependent effects that protect plants from UV-induced stress. The UV-A1 and UV-B irradiances used in the study were comparable to those in natural sunlight at the Earth surface making the observations relevant to plants growing in the field. The findings emphasize the role of photoreceptors in light perception, acclimation, and photoprotection of photosynthesis.

Analyses of photosynthesis, including quantum yield of PSII photochemistry (φPSII and Fv/Fm) and NPQ as well as biochemical tests, revealed a major role of CRYs in the maintenance of photosynthetic performance under exposure to UV-B, UV-A1 and blue radiation. We show in agreement with (Davey *et al*., 2012) that low levels of UV-B (1 µmol m^-2^ s^-1^) in the absence of UV-A and visible light had no effects on PSII photochemistry in the *uvr8* mutants (*uvr8-2* and *uvr8-7*) as compared to their respective wildtypes after 20 h of exposure. These findings may indicate that UVR8 alone is not required for optimal photosynthetic performance at low rates of UV-B. However, when UVR8 signaling was inactive together with impaired CRY activity, a large reduction in photoprotection was observed in *uvr8-2cry1cry2* under all treatments which was more pronounced under UV-B and UV-A1 in the presence of blue light (Fig. 1). Thus, functional redundance in maintaining photoprotection and photosynthetic performance under UV and blue light could be one of the mechanisms by which UVR8 and CRYs promote plant survival under natural conditions (Rai et al., 2019; Tissot and Ulm, 2020). Interestingly, overexpression of UVR8 showed better protection of PSII activity in all light treatments but clearly enhanced in plants exposed to UV-B in the presence of blue light (Fig. 2). It is possible that high flavonol contents in *UVR8-OE* (Fig. 8) together with other phenolic compounds (Leonardelli *et al*., 2024) provide better protection of PSII in this mutant.

Another factor that may differentially influence photosynthesis under the used light treatments is gas exchange through stomata. Blue light stimulates stomatal opening through phototropins and CRYs (Wang et al., 2020) while increases in aperture in response to UV-A1 (360 nm), UV-A1 + Blue (459 nm), UV-B (287 nm) and UV-B + Blue have been also reported for Arabidopsis epidermal peels, with UV-A1 + Blue eliciting much smaller apertures than any of UV-B, UV-A1 or Blue on their own (Eisinger et al. 2003). While these authors used lower UV irradiances than us, Tossi et al. (2014) also using epidermal peels, reported that stomatal opening in white light is partly reverted by added UV-B, but only at irradiances higher than those used here. However, at the irradiances used, differential stomatal opening in response to the treatments likely lead only to moderate differences in intercellular CO2 concentration unlikely to significantly limit carboxylation (c.f. Wang et al. 2020, who grew their plants in the same GR). The availability of CO2 for Calvin-Benson-Bassham cycle can modify light reactions of photosynthesis and kinetics of chlorophyll fluorescence. Different response of wildtype plants in Figures 1 and 2 to UV-A1 and UV-B alone versus UV supplemented with blue could be partially due to differences in stomatal opening. Importantly, however, the endpoint values of dark-adapted Fv/Fm were indistinguishable between UV and UV + blue treatments (Fig. 3), indicating that possible blue-light stomatal opening did not contribute to protection of photosynthetic apparatus from photoinhibition. Further studies are required to evaluate the impact of stomatal function on photosynthetic performance under combined spectral treatments.

Exposure to UV-A1 (100 µmol m^-2^ s^-1^) and UV-B (1 µmol m^-2^ s^-1^) was accompanied with dramatic changes in photosynthetic electron transport chain and its redox states, as assessed by LHCII phosphorylation. Phosphorylation level of LHCII depends on the redox state of thylakoid plastoquinone pool between PSII and PSI. As a rule, under non-saturating light intensities the more active is PSII relatively to PSI, the more reduced is the plastoquinone pool in illuminated leaves, and hence the more phosphorylated is LHCII. Accordingly, low levels of LHCII phosphorylation may be caused either by exposure to light spectra that are preferentially absorbed by PSI as compared to PSII (Mattila *et al*., 2020), or by suppressed photochemistry of PSII, e.g., due to qI. UV treatments performed in this study affected the redox states of photosynthetic electron transport chain in two clearly different ways. Under UV-B, LHCII phosphorylation was completely missing, while it was restored under UV-B plus blue. If this was due to qI, it would have been accompanied by lower Fv/Fm under UV-B than under UVB plus blue, which was not the case (Fig. 3). Thus, the absent LHCII phosphorylation under UV-B was not caused by PSII photoinhibition, rather by preferential absorbance of UV-B by PSI. This interpretation is supported by negative apparent NPQ values after 30 min of exposure to UV-B. The reason for negative NPQ is that Fm’ under UV-B was higher than dark-adapted Fm. This strongly suggests that UV-B promoted association of mobile pool of LHCII to PSII (formation of state 1), which increased light-absorption cross-section of PSII and thus Fm’ (Fig. S3). Taken together, these observations strongly indicate that the used UV-B treatment favoured PSI excitation and could thus be considered a “PSI light” (Mattila *et al*., 2020) that promoted photosynthetic state transitions (qT) to state 1.

In contrast to UV-B, UV-A1 treatment did not fully suppress LHCII phosphorylation in the wildtype L*er* (Fig. 6). This indicated that the defects in LHCII phosphorylation observed in the mutants under UV-A1 were most probably due to suppressed photochemistry of PSII. This agrees with chlorophyll fluorescence analyses (Figs 1, 3, 4). In the absence of functional CRYs, LHCII phosphorylation was strongly impaired by UV-A1, while in the absence of UVR8 and CRYs it was entirely abolished under UV-A1. Thus, both UVR8 and CRYs together were required for proper function of PSII, specifically under high UV-A1 irradiance. Overall, the results supported important and partially redundant roles of CRYs and UVR8 in acclimation of plant photosynthetic apparatus to UV. The effects of several UV-A wavebands on redox states of plastoquinone pool have been reported (Mattila *et al*., 2020). Our observations extend the knowledge to more short-wave spectral regions including UV-B. This sets the basis for further studies of the roles of UV and photoreceptors in photosynthetic light harvesting and in energy distribution between PSII and PSI.

Our data corroborated the importance of UVR8 in mediating the accumulation of epidermal flavonols in plants exposed to UV-B and UV-A radiation (Favory *et al*., 2009; Morales *et al*., 2013; Rai *et al*., 2019) and that of CRYs under blue light and UV-A radiation (Rai *et al*., 2019). Both *uvr8-2* and *uvr8-7* accumulated flavonols to lower levels than their respective wildtypes in plants exposed to UV-B (Figs 7, 8). Furthermore, impaired UVR8 and CRY signaling in *uvr8-2cry1cry2* resulted in lower flavonol levels under all light treatments. The accumulation of epidermal flavonols was also lower in *cry1cry2* than in L*er*. The resulting weaker epidermal attenuation of incoming UV radiation can be expected to provide only partial protection against photoinhibition (Figs 3, 7). In agreement with this, enhanced levels of NPQ, most likely its qE component, in *cry1cry2* and *uvr8-2cry1cry2* under short-term UV-A1 or UV-A1 plus blue treatments suggested that the photosynthetic apparatus of these mutants was more exposed to UV-A1 than in other genotypes. This mechanism corresponds to the first “line of defence”, or reduced exposure of chloroplasts.

The low flavonol levels measured with Dualex in these experiments correspond also to low kaempferol and quercetin levels determined in L*er*, *uvr8-2*, *cry1cry2* and *uvr8-2cry1cry2* under simulated UV-B, UV-A and blue (Rai et al 2019). Thus, it is possible that these flavonols, together with other phenolics (Leonardelli *et al*., 2024), could play key roles in protecting plants from photoinhibition under the light conditions tested. Our data show contrasting patterns of flavonol and anthocyanin accumulation in *uvr8-2cry1cry2* under all light treatments except UV-B in the presence of blue light (Figs. 7, S7). Also, *UVR8*-OE showed higher flavonol levels but lower anthocyanin content under initial GR conditions and UV-B exposure than Ws (Figs. 8, S9). These findings collectively suggest that anthocyanin may not be as important for photoprotection as flavanols under our experimental conditions and that UVR8 favoured the synthesis of flavonols in part at the expense of anthocyanins under UV-B. Interestingly, overexpression of UVR8 also resulted in better tolerance to MV. The high levels of protection in *UVR8*-OE seems to be dependent at least in part on more efficient scavenging of ROS than in the wildtype. This mechanism could correspond to the second line of defence, i.e., dissipation of excess energy and prevention of oxidative damage.

In conclusion, as efficient photosynthetic carbon fixation and mechanisms for the prevention of damage are in conflict, a delicate balance is maintained and adjusted through multiple mechanisms. Some of this regulation is achieved through feedback within the photosynthetic regulatory mechanisms. Here we show that UVR8 and CRY dependent signaling also plays a crucial role in this regulation. Our data with different photoreceptor mutants highlight the importance of photoreceptor activity in the context of interaction with other photoreceptors. The results presented here expand our understanding of the links between light perception and photoprotection with broad implications for plant performance in natural and managed environments.

## Acknowledgements

LOM acknowledges the Swedish Research Council FORMAS (https://formas.se/en; grant #2021-00616) and the Magnus Bergvalls foundation. MB was supported by grants from Research Council of Finland Centre of Excellence in Molecular Biology of Primary Producers (2014–2019, decision 271832 and 307335), and 349540. AS was supported by the Centre of Excellence in Tree Biology, Research Council of Finland (decision 346140).

## Author contributions

LOM, AS and MB conceived and designed the research. LOM, AS and MB conducted photosynthesis experiments and data analysis. PJA assembled LEDs and designed the light treatments. NR performed Dualex measurements and data analysis. LOM, AS and MB wrote the manuscript with contributions by PJA and NR.

