## Supplemental data for "Protection of photosynthesis by UVR8 and cryptochromes in Arabidopsis under blue and UV radiation"

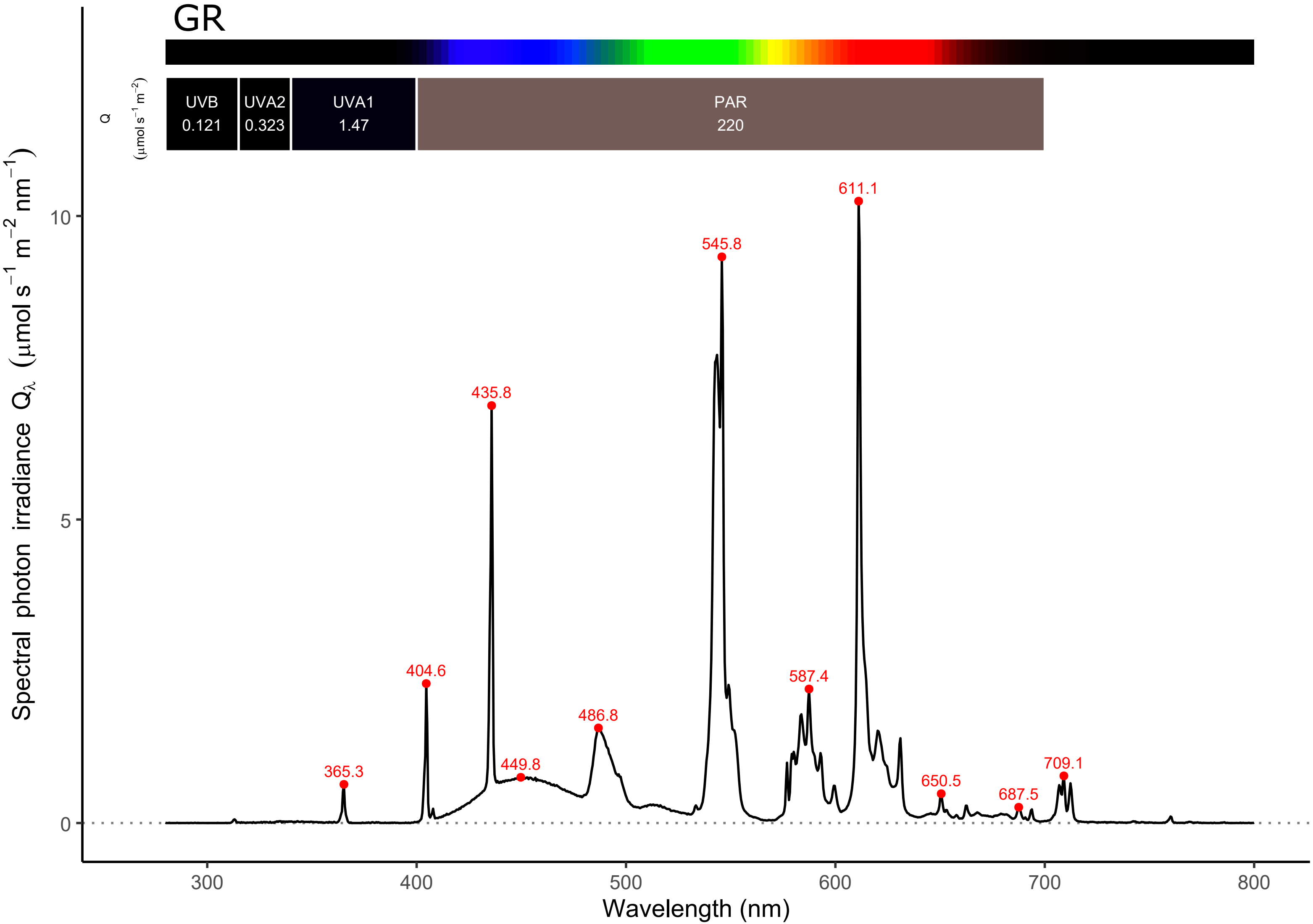

GR
UVB
0.121
UVA2
0.323
UVA1
1.47
PAR
220
Q
(μmol s⁻¹ m⁻²)
611.1
545.8
435.8
404.6
587.4
486.8
365.3
449.8
709.1
650.5
687.5
10
5
0
300
400
500
600
700
800
Wavelength (nm)
Spectral photon irradiance Qλ (μmol s⁻¹ m⁻² nm⁻¹)

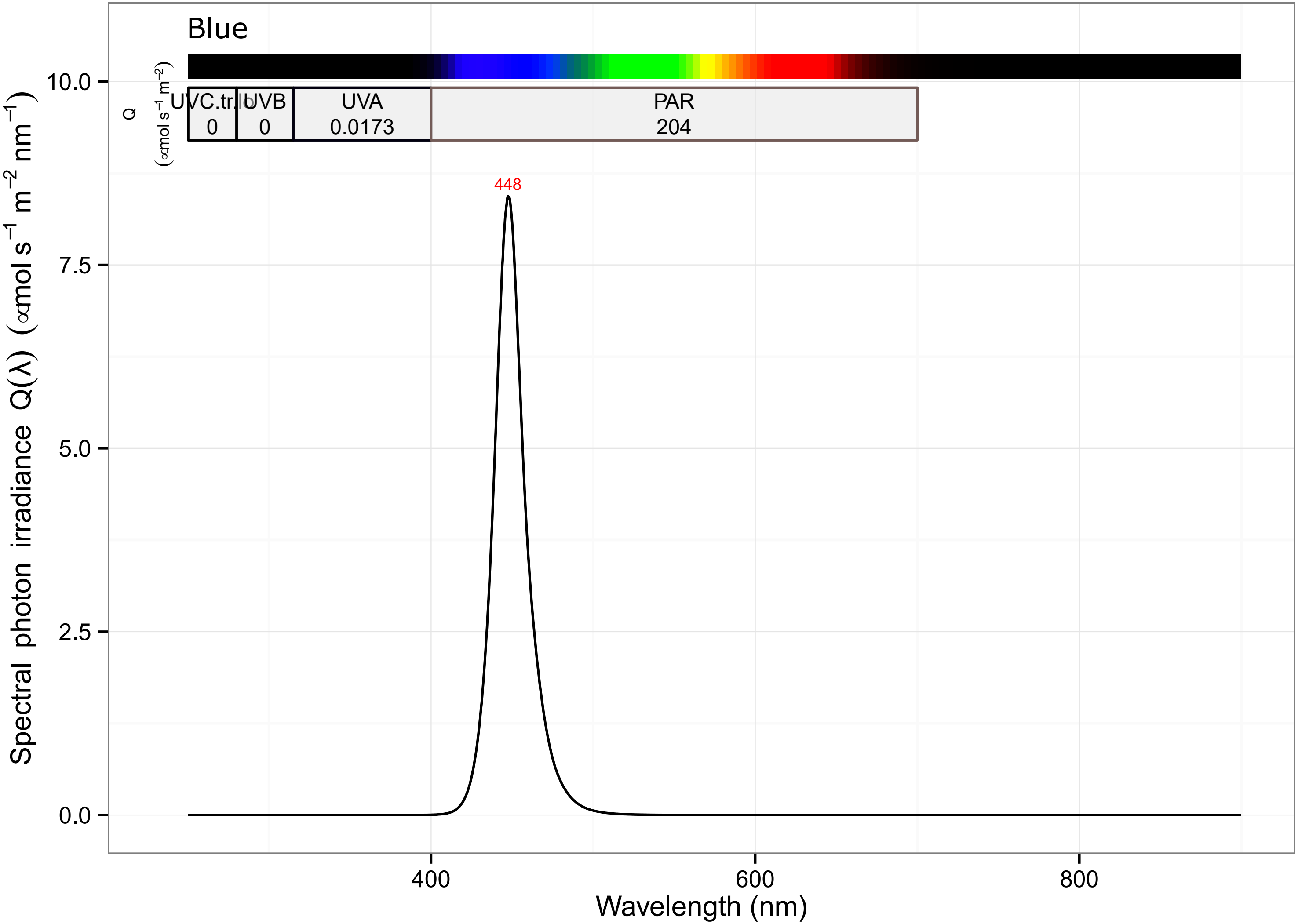

Blue
UVC.tr.lo
0
UVB
0
UVA
0.0173
PAR
204
Q
(∝mol s⁻¹ m⁻²)
448
10.0
7.5
5.0
2.5
0.0
400
600
800
Wavelength (nm)
Spectral photon irradiance Q(λ) (∝mol s⁻¹ m⁻² nm⁻¹)

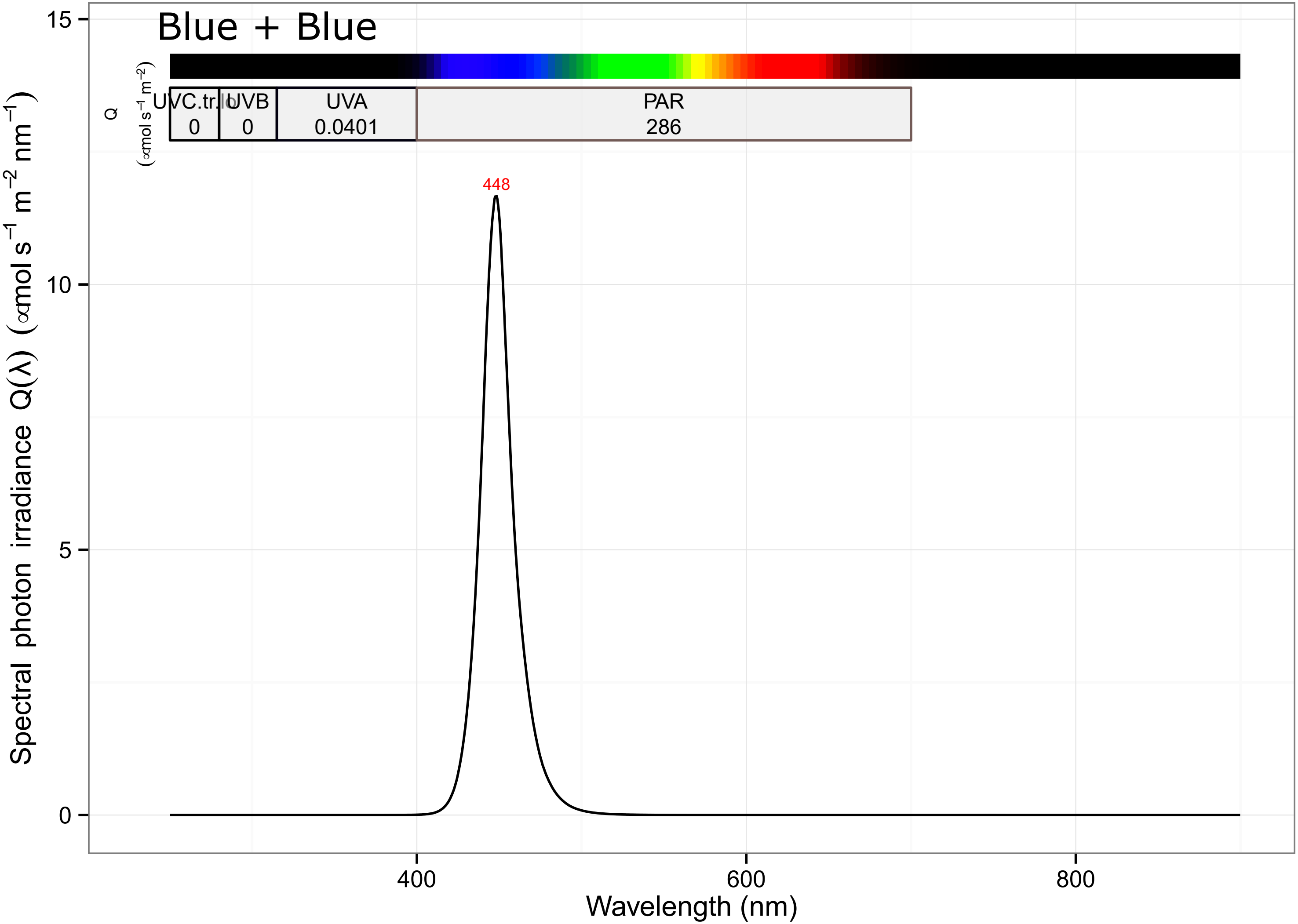

Blue + Blue
UVC.tr.lo
0
UVB
0
UVA
0.0401
PAR
286
Q
(µmol s⁻¹ m⁻²)
448
Spectral photon irradiance Q(λ) (µmol s⁻¹ m⁻² nm⁻¹)
15
10
5
0
400
600
800
Wavelength (nm)

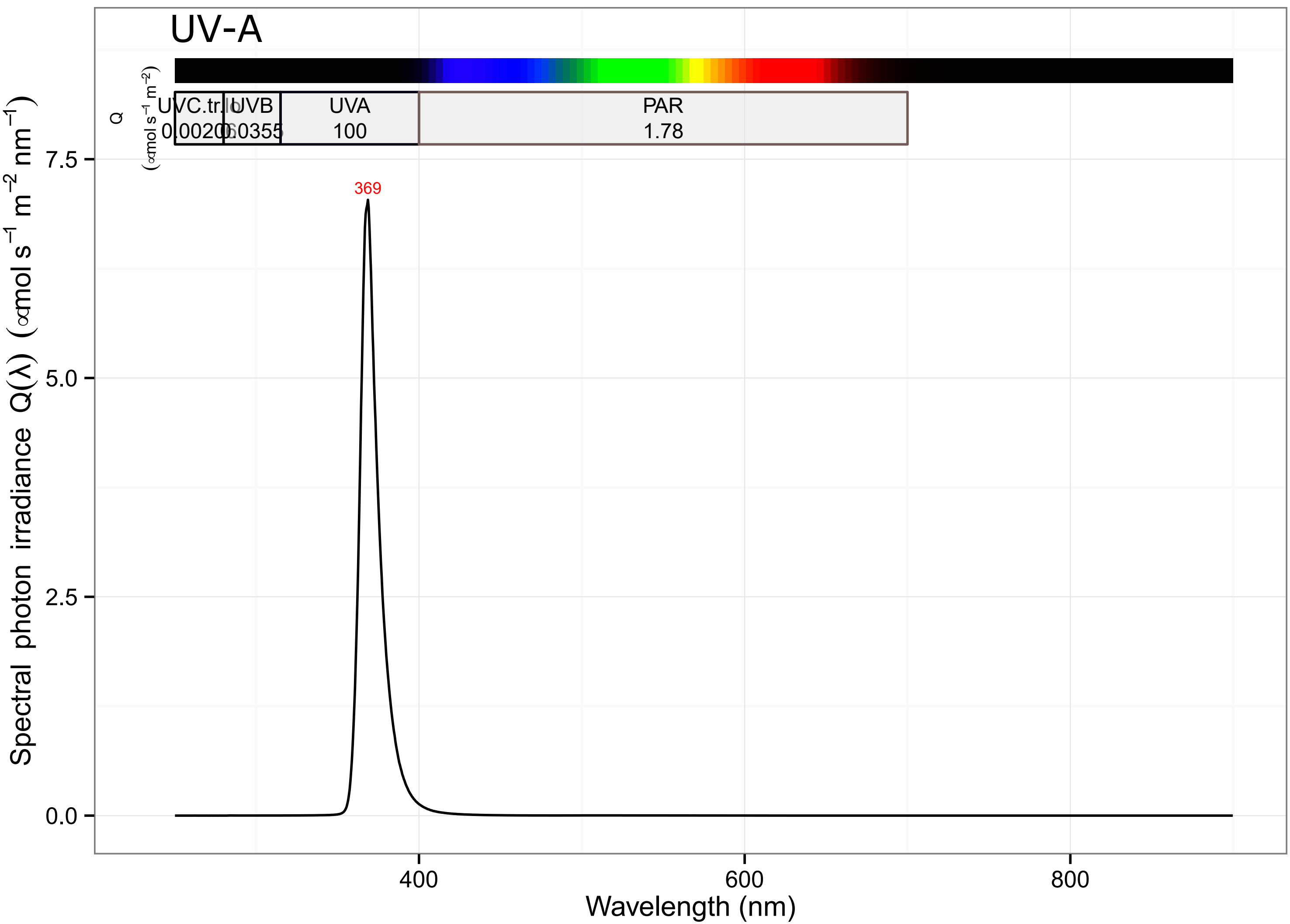

UV-A
UVC.tr.lo
UVB
UVA
PAR
0.0020
0.0355
100
1.78
Q
(∝mol $s^{-1}$ $m^{-2}$)
369
7.5
5.0
2.5
0.0
Spectral photon irradiance Q(λ) (∝mol $s^{-1}$ $m^{-2}$ $nm^{-1}$)
400
600
800
Wavelength (nm)

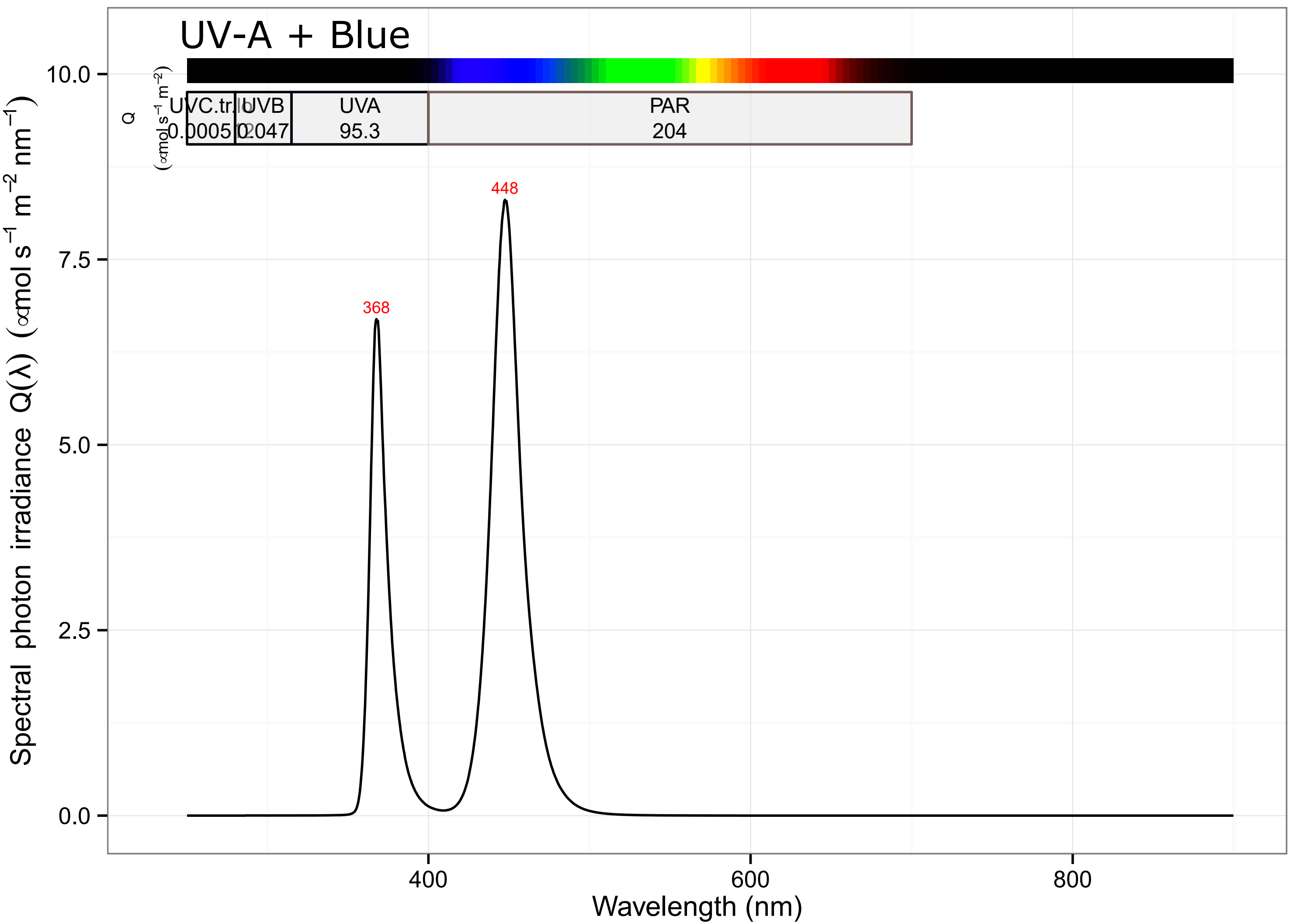

UV-A + Blue
UVC.tr.lo
0.0005
UVB
0.2047
UVA
95.3
PAR
204
Q
(∝mol s⁻¹ m⁻²)
448
368
10.0
7.5
5.0
2.5
0.0
400
600
800
Wavelength (nm)
Spectral photon irradiance Q(λ) (∝mol s⁻¹ m⁻² nm⁻¹)

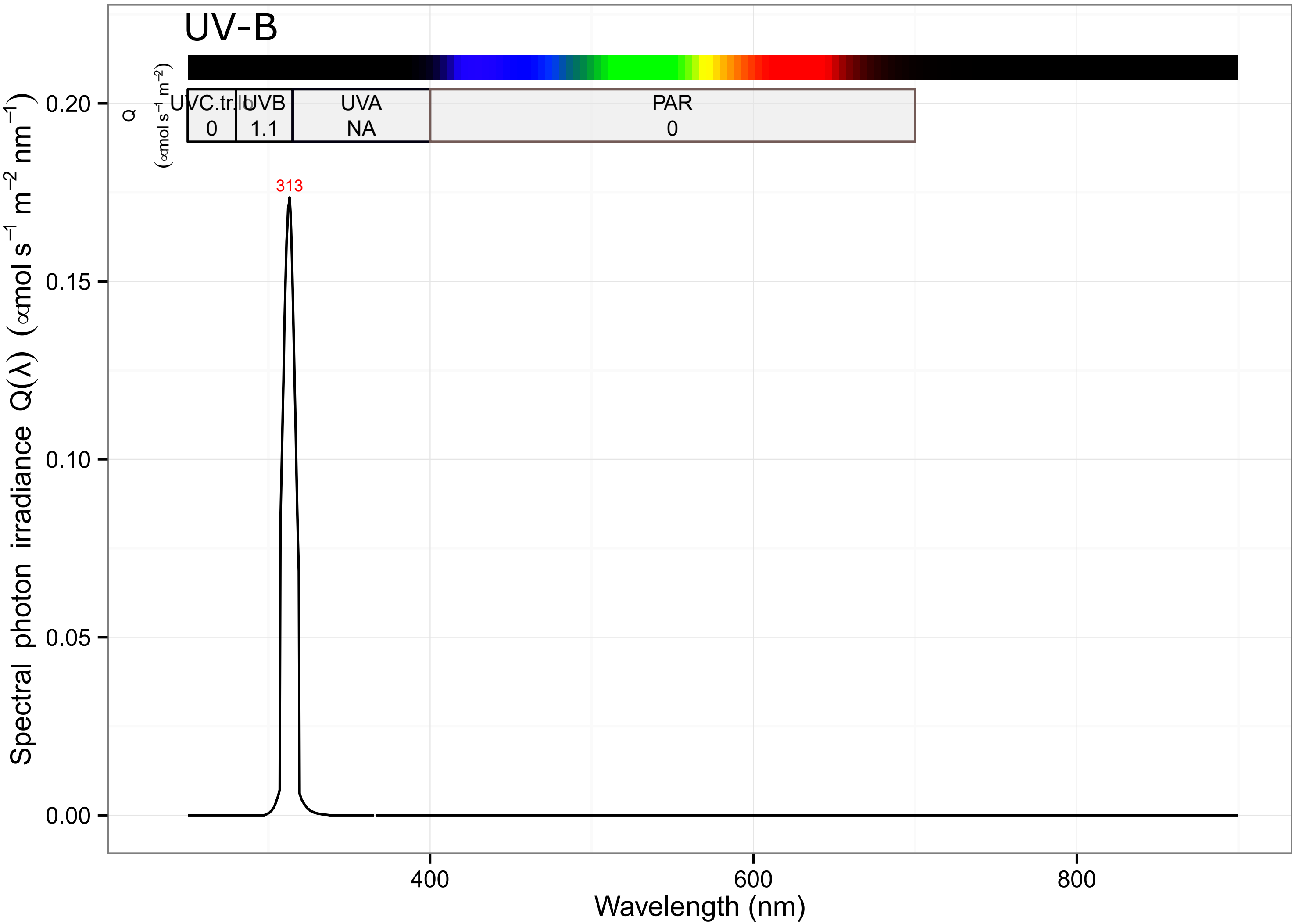

UV-B
UVC.tr.lo
0
UVB
1.1
UVA
NA
PAR
0
Q
(∝mol s⁻¹ m⁻²)
313
0.20
0.15
0.10
0.05
0.00
Spectral photon irradiance Q(λ) (∝mol s⁻¹ m⁻² nm⁻¹)
400
600
800
Wavelength (nm)

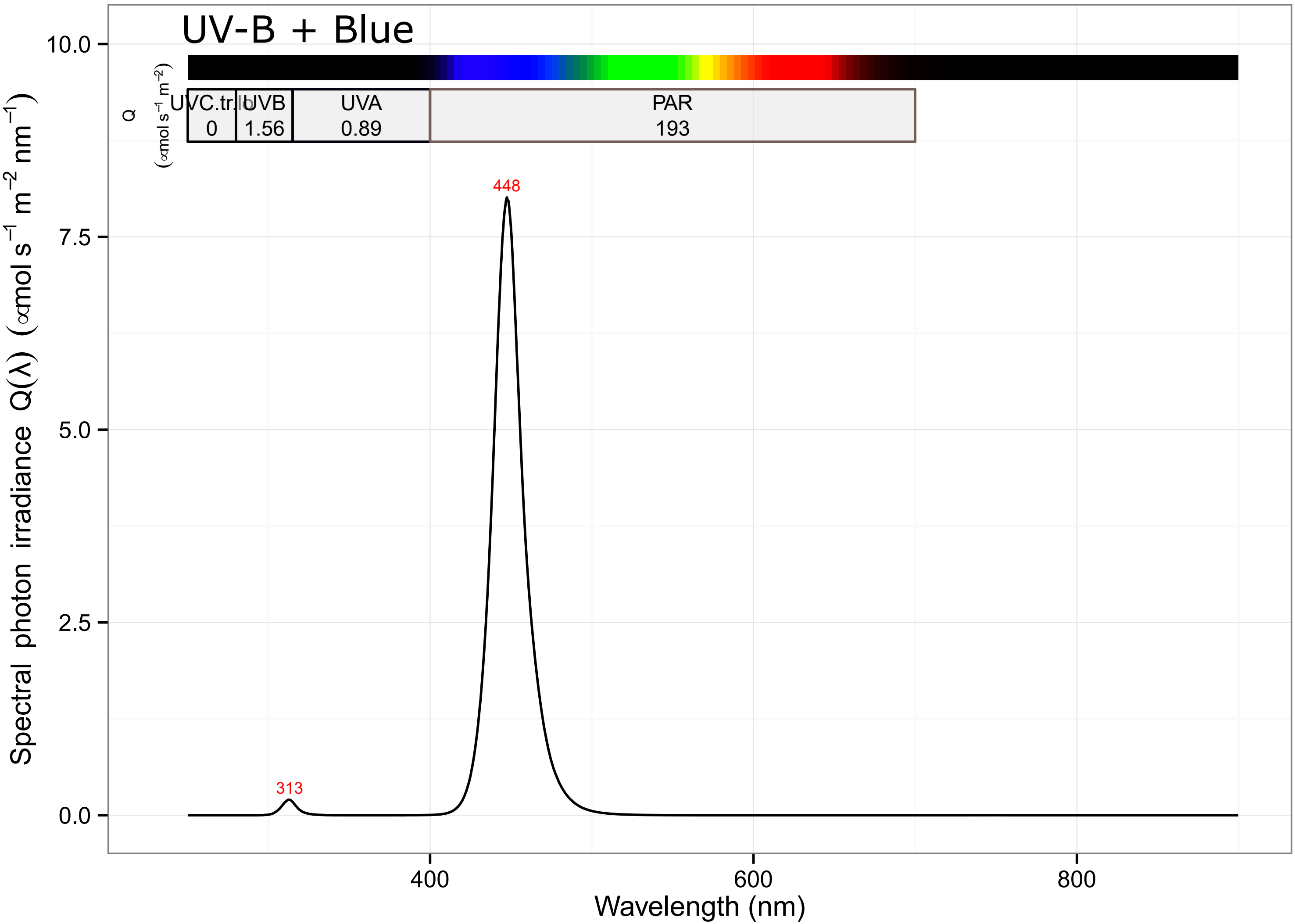

UV-B + Blue
Q
(∝mol s$^{-1}$ m$^{-2}$)
UVC.tr.lo
0
UVB
1.56
UVA
0.89
PAR
193
448
313
10.0
7.5
5.0
2.5
0.0
400
600
800
Spectral photon irradiance Q(λ) (∝mol s$^{-1}$ m$^{-2}$ nm$^{-1}$)
Wavelength (nm)

### Fig. S2

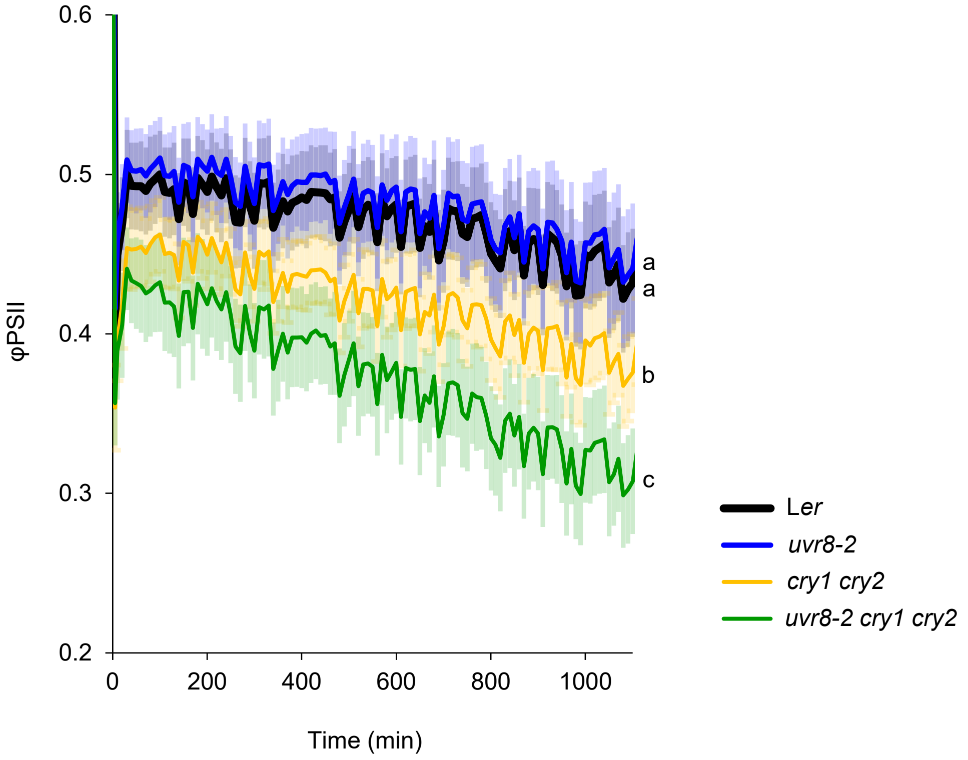

0.6
0.5
0.4
0.3
0.2
φPSII
0
200
400
600
800
1000
Time (min)
a
a
b
c
Ler
uvr8-2
cry1 cry2
uvr8-2 cry1 cry2

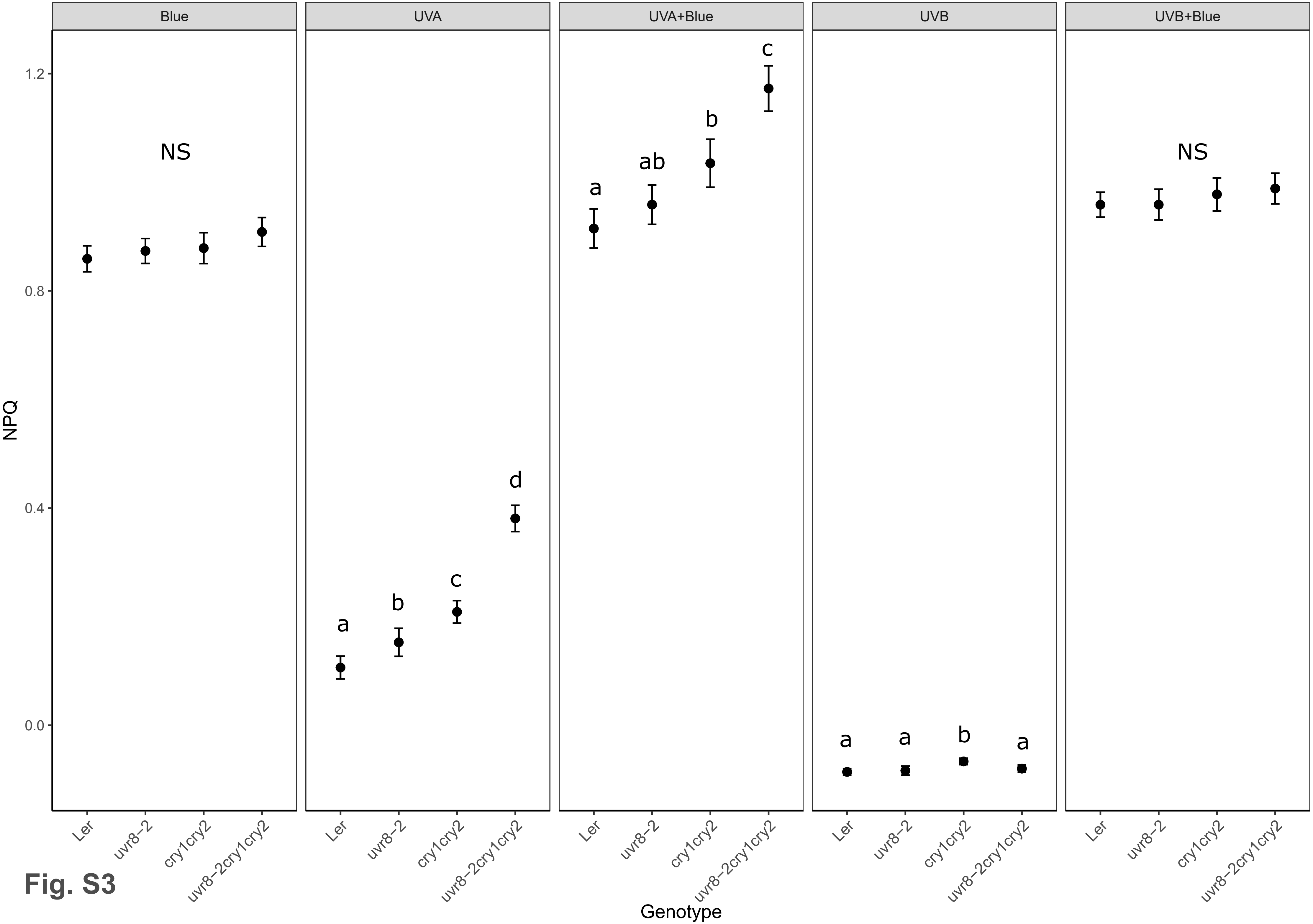
Blue
UVA
UVA+Blue
UVB
UVB+Blue
NPQ
1.2
0.8
0.4
0.0
NS
a
b
c
d
a
ab
b
c
a
a
b
a
NS
Ler
uvr8−2
cry1cry2
uvr8−2cry1cry2
Genotype

**Fig. S3**

### Fig. S4

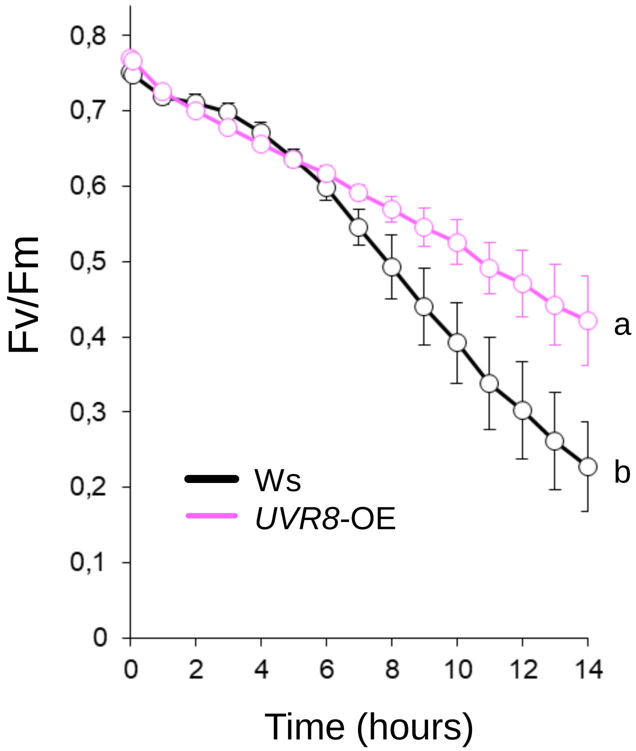

Fv/Fm
0,8
0,7
0,6
0,5
0,4
0,3
0,2
0,1
0
0
2
4
6
8
10
12
14
Time (hours)
Ws
UVR8-OE
a
b

### Fig. S5

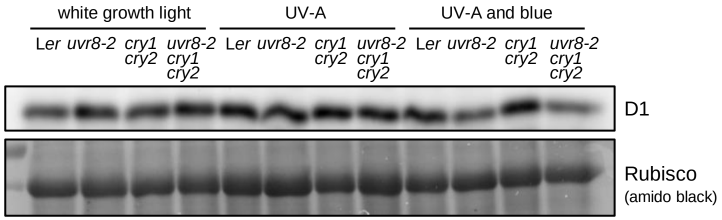

white growth light
UV-A
UV-A and blue
Ler uvr8-2 cry1 cry2 uvr8-2 cry1 cry2 Ler uvr8-2 cry1 cry2 uvr8-2 cry1 cry2 Ler uvr8-2 cry1 cry2 uvr8-2 cry1 cry2
D1
Rubisco
(amido black)

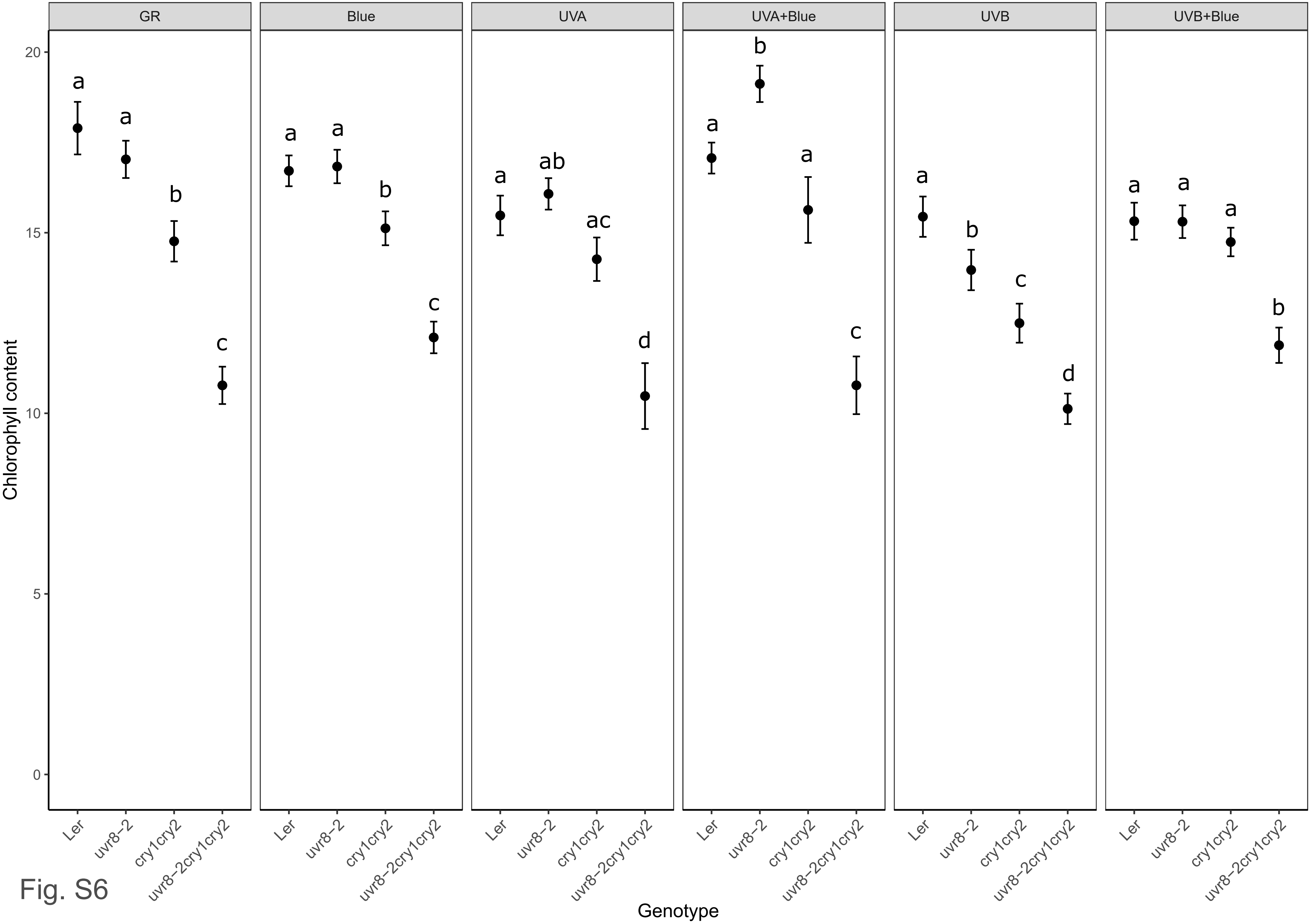

GR
Blue
UVA
UVA+Blue
UVB
UVB+Blue
Chlorophyll content
0
5
10
15
20
a
a
b
c
a
a
b
c
a
ab
ac
d
a
b
a
c
a
b
c
d
a
a
a
b
Ler
uvr8−2
cry1cry2
uvr8−2cry1cry2
Genotype


Fig. S6

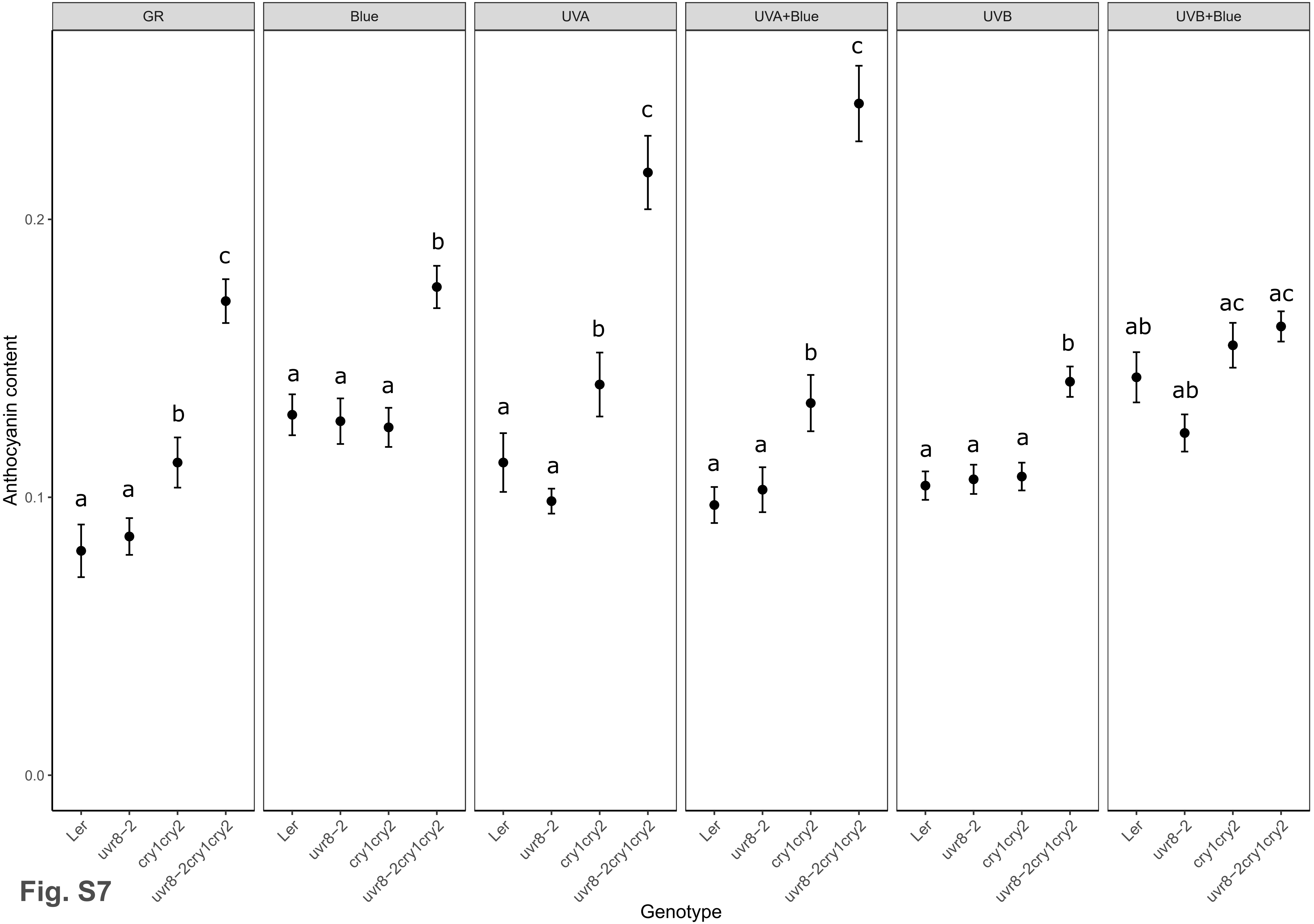
GR
Blue
UVA
UVA+Blue
UVB
UVB+Blue
Anthocyanin content
0.0
0.1
0.2
a
a
b
c
a
a
a
b
a
a
b
c
a
a
b
c
a
a
a
b
ab
ab
ac
ac
Ler
uvr8−2
cry1cry2
uvr8−2cry1cry2
Genotype

**Fig. S7**

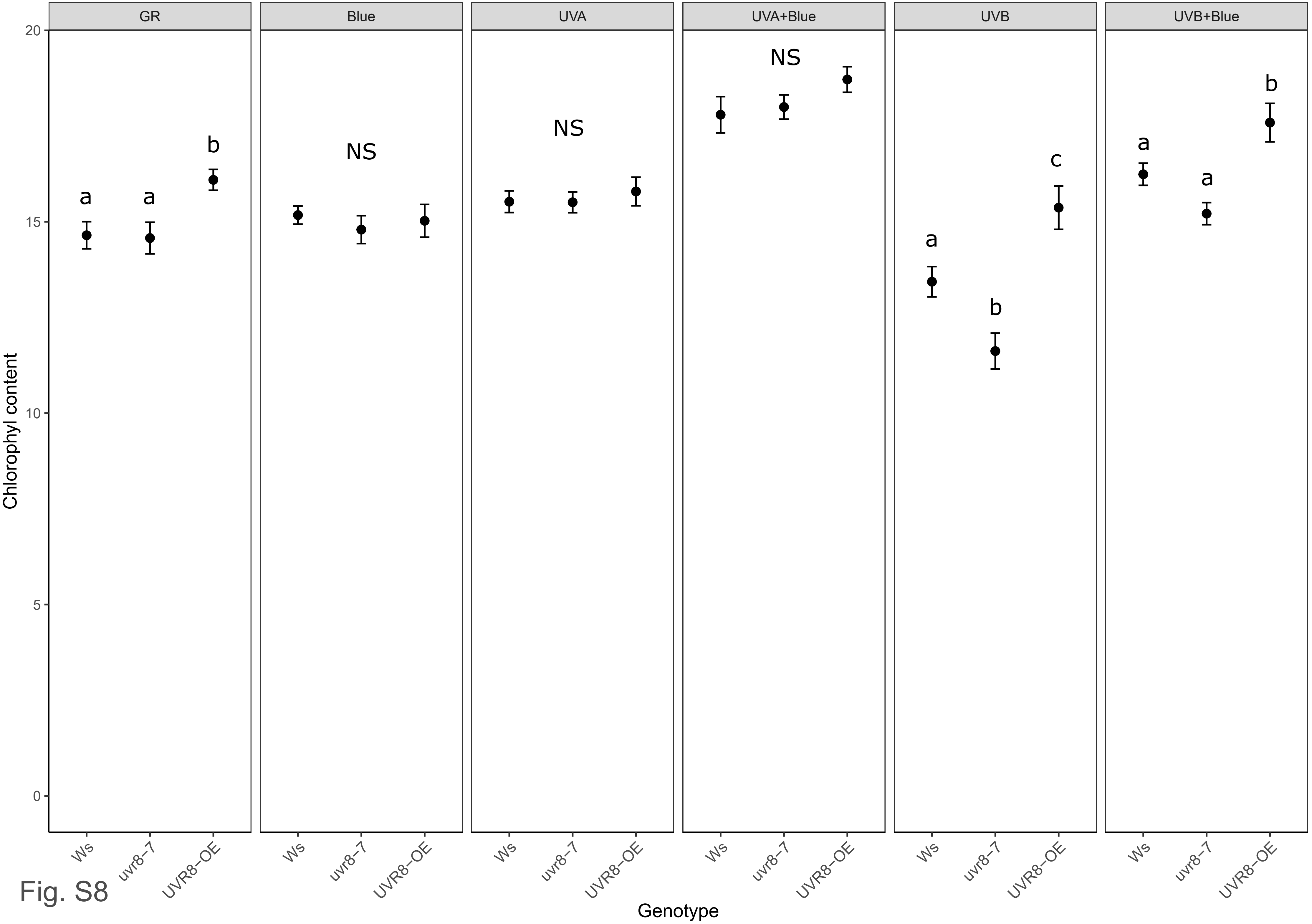
GR
Blue
UVA
UVA+Blue
UVB
UVB+Blue
Chlorophyl content
20
15
10
5
0
a
a
b
NS
NS
NS
a
b
c
a
a
b
Ws
uvr8–7
UVR8–OE
Genotype
Fig. S8

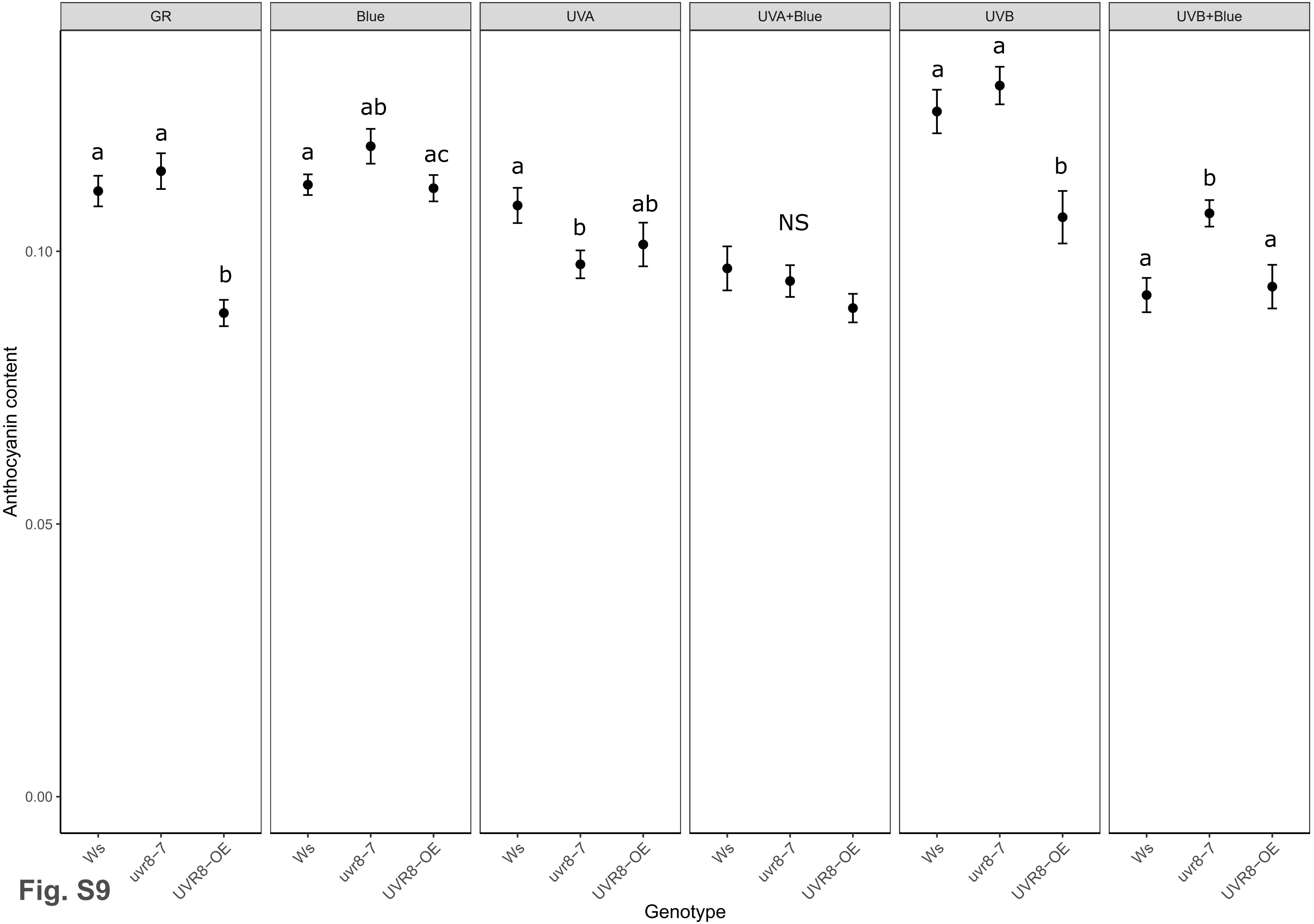
GR
Blue
UVA
UVA+Blue
UVB
UVB+Blue
a
a
b
a
ab
ac
a
b
ab
NS
a
a
b
b
a
a
Anthocyanin content
0.10
0.05
0.00
Ws
uvr8−7
UVR8−OE
Genotype
Fig. S9
